# Structural modeling reveals the mechanism of motor ATPase coordination during type IV pilus retraction

**DOI:** 10.1101/2025.10.30.685630

**Authors:** Abigail E. Teipen, Jacob D. Holt, Diane L. Lynch, Yixuan Peng, Triana N. Dalia, James C. Gumbart, Carey D. Nadell, Ankur B. Dalia

## Abstract

Diverse bacterial species utilize surface appendages called type IV pili (T4P) to interact with their environment. These structures are dynamically extended and retracted from the cell surface, which is critical for diverse functions. Some T4P systems rely on two distinct motor ATPases, PilT and PilU, whose combined activities are required to power forceful T4P retraction. However, the mechanism by which these motors coordinate to facilitate T4P retraction has remained unclear. Here, we utilize the competence T4P in *V. cholerae* as a model system to elucidate the molecular basis for PilT-PilU coordination during T4P retraction. Specifically, we modeled the interactions between PilT and PilU using AlphaFold 3 and molecular dynamics (MD) simulations. We then empirically tested these models using a combination of cytological and high-resolution genetic approaches. Our results reveal that interactions between PilT and the PilU C-terminus are critical for these motors to coordinate to drive T4P retraction. Finally, we show that PilT-PilU interactions are broadly conserved in T4P systems from diverse bacterial species, and we experimentally validate that they are required for T4P retraction in *Acinetobacter baylyi*. Together, this work expands our fundamental understanding of T4P dynamics, and more broadly it provides mechanistic insight into how these ATPases coordinate to assemble some of the strongest biological motors in nature.

**SIGNIFICANCE:** Diverse bacterial species use filamentous surface appendages called type IV pili (T4P) to move along surfaces, take up DNA for horizontal gene transfer, and stick to biotic and abiotic surfaces. The forceful retraction of these filaments is often required for these behaviors. In many T4P systems, the combined activity of two distinct motor ATPase proteins is required for forceful retraction; however, a detailed understanding of *how* these motor proteins interact to promote forceful retraction is currently lacking. Here, we use an integrated approach to uncover the molecular mechanism for motor ATPase coordination. Furthermore, we show that this mechanism is broadly conserved in diverse T4P systems.

## INTRODUCTION

Type IV pili (T4P) are broadly conserved proteinaceous surface appendages that mediate several important behaviors including DNA uptake for natural transformation, surface attachment, surface sensing, and twitching motility (1–5). The T4P machine is a large macromolecular complex composed of at least 80 proteins (via a multimeric assembly of 7 unique proteins) (6–8). This nanomachine dynamically assembles (extends) and disassembles (retracts) a pilus fiber through an outer membrane gated secretin (9, 10), and this dynamic extension and retraction is critical for the diverse functions carried out by T4P (2, 4, 11).

In canonical T4P (*i.e.*, type IVa pilus systems), dynamic extension and retraction are powered by distinct hexameric motor ATPases. Pilus extension is often driven by a single motor ATPase protein, PilB, that interacts with the T4P platform protein, PilC, and hydrolyzes ATP to power the assembly of major pilin subunits into a helical pilus filament that extends from the cell surface through the outer membrane secretin, PilQ (12, 13). Conversely, T4P retraction is commonly powered by two distinct motor ATPases – PilT and PilU. Previous studies have established that PilT serves as the primary retraction ATPase (12, 14, 15) that directly interacts with PilC to promote depolymerization of major pilin subunits. PilU is an accessory ATPase (16, 17) that drives forceful T4P retraction in a PilT-dependent manner (18–20). A detailed mechanistic understanding of *why* PilU requires PilT to promote retraction and *how* PilT and PilU coordinate to promote forceful T4P retraction, however, remains unclear.

In this study, we use the *Vibrio cholerae* competence T4P as a model system to dissect the molecular mechanism underlying PilT-PilU-mediated retraction. The *V. cholerae* competence T4P is required for DNA uptake during horizontal gene transfer by natural transformation (NT) (1). During this process, competence T4P extend from the bacterial surface, bind to DNA in the environment, and then retract to bring this DNA into the cell (11, 21). This DNA is then translocated into the cytoplasm and if it is homologous to the host genome, it can be integrated by homologous recombination (22). NT is induced when *V. cholerae* colonizes the chitinous shells of crustacean zooplankton in the aquatic environment (23). Recent work highlights that PilU-dependent retraction is necessary for NT when DNA is adsorbed to a chitin surface (24). Thus, PilT and PilU are both critical for facilitating DNA uptake by competence T4P under physiologically relevant conditions. Also, the *V. cholerae* competence T4P has emerged as an invaluable model system for studying the basic biology of T4P (11, 18, 19, 25–27). Thus, it is ideally suited for dissecting the molecular mechanism underlying PilT-PilU coordination during T4P retraction.

## RESULTS

### AlphaFold 3 models interactions between the retraction motor ATPases and the T4P machine

To define how PilT and PilU coordinate to facilitate T4P retraction, we first sought to model their interactions with PilC using AlphaFold 3 (AF3) (28). Due to the large size of this complex, we generated a hybrid model of PilC-PilT-PilU by independently modeling the interaction between 1) PilC and the PilT hexamer and 2) the PilT and PilU hexamers (**Fig. S1a-b**). Then, one subunit of PilT was aligned between these two models to generate the hybrid complex (**Fig. 1a**).

**Fig. 1.**
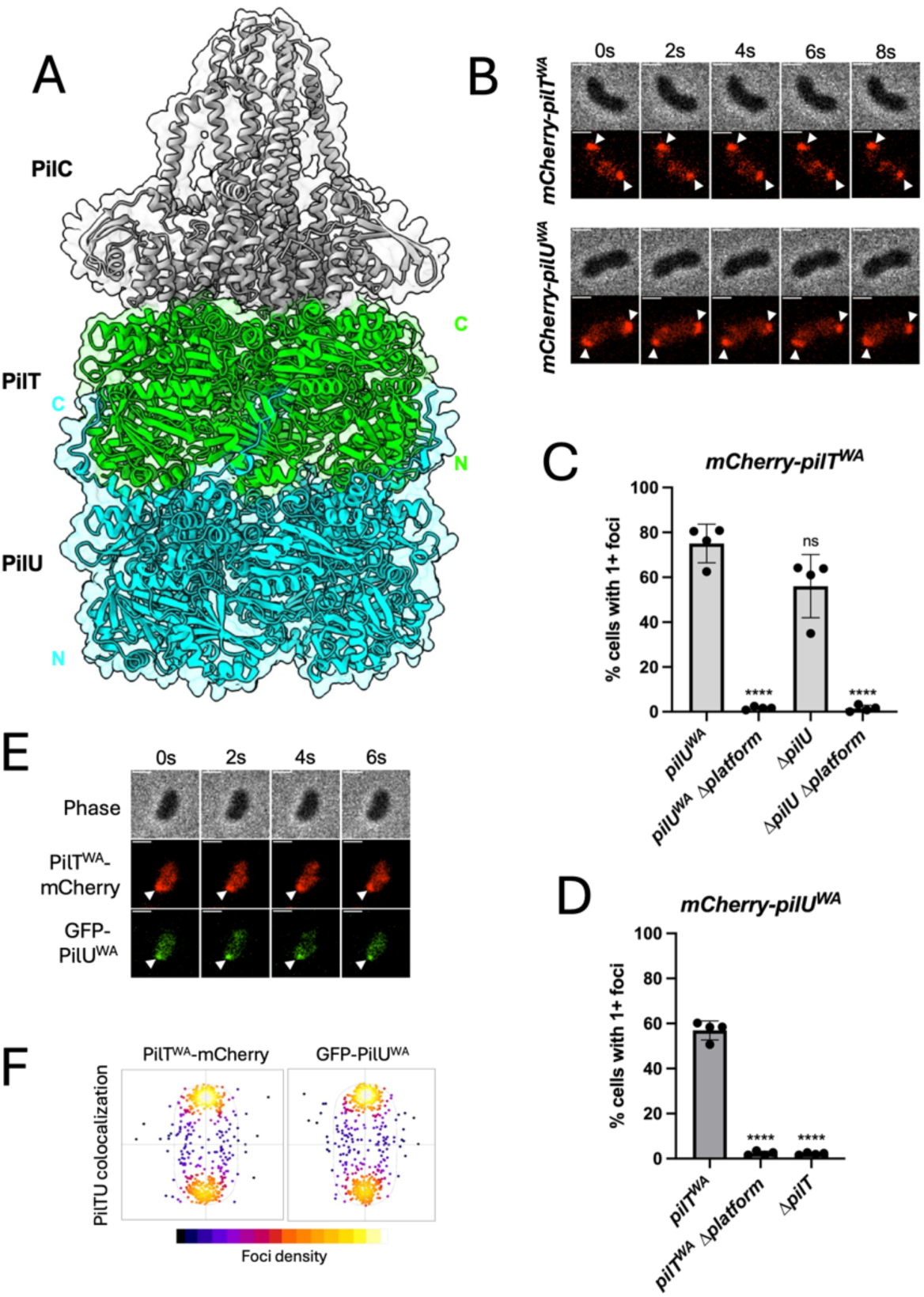
PilT bridges PilU to the T4P machine. (**A**) A hybrid AlphaFold 3 model highlights the putative interactions between PilC (grey), PilT (green), and PilU (cyan). The N-terminal and C-terminal faces of PilT and PilU are labeled with green and cyan letters, respectively. Motor localization assays were performed to assess the localization of PilT and PilU *in vivo* by epifluorescence time lapse microscopy in **B**-**D**. (**B**) Representative montages of timelapse imaging of *V. cholerae* cells expressing mCherry-PilT^WA^ PilU^WA^ and PilT^WA^ mCherry-PilU^WA^ that displayed ATPase foci (white carats). Scale bar, 1 µm. For quantification of these assays, a fluorescent focus was only counted if it was maintained throughout the duration of the 10-s timelapse. (**C**) Motor localization assay of cells expressing *mCherry-pilT^WA^* (WA = the *pilT^K136A^*Walker A mutation that ablates ATPase activity) and the indicated mutations. The Δ*platform* mutation denotes a mutant lacking both the competence T4P platform *pilC* and the MSHA T4P platform *mshG*. For each strain, the percentage of cells (*n* = 120 cells analyzed) with one or more fluorescent mCherry-PilT^WA^ foci is reported. (**D**) Motor localization assay of cells expressing *mCherry-pilU^WA^*(WA = the *pilU^K134A^* Walker A mutation that ablates ATPase activity) and the indicated mutations. For each strain, the percentage of cells (*n* = 120 cells analyzed) with one or more fluorescent mCherry-PilU^WA^ foci is reported. (**E**) Motor localization assay of cells expressing *pilT^WA^- mCherry gfp-pilU^WA^*. The representative montage of timelapse imaging shows colocalization of PilT^WA^-mCherry and GFP-PilU^WA^ (white carat). (**F**) Cell scatter plots show the localization of PilT^WA^-mCherry and GFP-PilU^WA^ foci in the colocalization strain shown in **E**. Both PilT and PilU foci primarily localize to the cell poles. Data in **C**-**D** are from 4 independent biological replicates and shown as the mean ± SD. Statistical comparisons were made by one-way ANOVA with Tukey’s multiple comparison test of the log-transformed data. ns, not significant; **** = *p* < 0.0001. Symbols directly above bars denote comparisons to the appropriate parent strain.

The relatively high model confidence scores for PilT-PilC (ipTM = 0.66) and PilT-PilU (ipTM = 0.63) suggest that PilT may directly interact with both PilC and PilU (**Fig. S1a-b**). By contrast, AF3 modeling of PilU-PilC yields very low confidence models (ipTM = 0.37), which suggests that PilU and PilC may not directly interact (**Fig. S1c**). Because PilT is predicted to interact with PilC and PilU via distinct interfaces (**Fig. 1a**), we hypothesize that PilU requires PilT to engage the T4P machine. This model is consistent with prior work demonstrating that PilU can only facilitate T4P retraction in the presence of PilT (18, 19, 26).

### PilT bridges PilU to the T4P machine

To test whether PilU requires PilT to engage the membrane-embedded T4P machine, we sought to develop an assay to examine the interaction of these ATPases with the T4P machine in live cells. When the ATPase activity of these motors is intact (*pilT^+^* and/or *pilU^+^*), it is likely that they only engage the T4P machine transiently while actively promoting T4P retraction. Accordingly, strains expressing *mCherry-pilT^+^* and *mCherry-pilU^+^* do not form foci at the cell periphery (**Fig. S2, Movies S1 and S2**). We hypothesized that we could stabilize the interaction of these ATPases with the T4P machine by ablating their ATPase activity via mutation of the <u>W</u>alker <u>A</u> (WA) site required for ATP-binding (*pilT^WA^* = *pilT^K136A^*; *pilU^WA^* = *pilU^K134A^*) (18, 29) (**Fig. S1b**). Indeed, in strains expressing *mCherry-pilT^WA^ pilU^WA^* and *pilT^WA^ mCherry-pilU^WA^*, fluorescent foci are readily observed at the cell periphery in the majority of cells (**Fig. 1b-d, Fig. S2c, Movies S3 and S4**). Also, these peripheral foci exhibited low mobility (**Fig. S2b**), which is consistent with these peripheral foci representing T4P machine-engaged motor ATPases. PilT and PilU can promote retraction of both the competence T4P and the MSHA T4P in *V. cholerae* (19, 30, 31) via interactions with the platform protein of each system (*pilC* and *mshG*, respectively). In a background where these platform proteins are deleted, mCherry-PilT^WA^ and mCherry-PilU^WA^ foci are ablated (**Fig. 1c-d**). This supports the hypothesis that the peripheral *mCherry-pilT^WA^* and *mCherry-pilU^WA^* foci observed in the parent represent T4P machine-engaged motors. To further test this hypothesis, we assessed whether these motor foci colocalize with the T4P machine. Specifically, we assessed the colocalization of mCherry-PilT^WA^ and mCherry-PilU^WA^ with competence T4P filaments and a competence T4P machine component (*msfGFP-pilQ*) in a background where the MSHA locus has been deleted (*ΔMSHA*), which ensures that motors can only interact with competence T4P machines. Qualitatively, we found that mCherry-PilT^WA^ and mCherry-PilU^WA^ foci colocalized with the base of extended competence T4P (**Fig. S3a, Movies S5 and S6**). Also, more quantitatively, we determined that mCherry-PilT^WA^ and mCherry-PilU^WA^ foci colocalize with msfGFP-PilQ foci at a frequency of 99.3 ± 0.7% and 98.1% ± 1%, respectively (**Fig. S3b-c, Movies S7 and S8**). All together, these results strongly suggest that mCherry-PilT^WA^ and mCherry-PilU^WA^ foci represent motor ATPases that are bound to a T4P machine.

The AF3 model suggests that PilT directly interacts with the T4P machine; thus, we hypothesized that PilU would be dispensable for PilT-T4P interactions. Consistent with this, *mCherry-pilT^WA^* Δ*pilU* cells still readily formed fluorescent foci (**Fig. 1c**). Importantly, deletion of the T4P platform proteins still ablates focus formation in this background (**Fig. 1c**). By contrast, the AF3 model suggests that PilT is required to bridge the interaction between PilU and the T4P machine (**Fig. 1a**). Indeed, fluorescent foci are ablated in *mCherry-pilU^WA^* Δ*pilT* compared to the *mCherry- pilU^WA^ pilT^WA^* parent (**Fig. 1d**). Together these results indicate that PilT may directly interact with the T4P machine, and that PilT is required to bridge PilU-T4P interactions.

The AF3 model also predicts a direct interaction between PilT and PilU. Accordingly, in *pilT^WA^- mCherry gfp-pilU^WA^* cells, 96% ± 2% of the GFP-PilU^WA^ foci overlap with a PilT^WA^-mCherry focus (**Fig. 1e, Movie S9**). Additionally, both PilT^WA^-mCherry and GFP-PilU^WA^ foci primarily localize to the cell poles (**Fig. 1f**) where T4P machines are most prevalent (**Fig. S3b-c**). This colocalization is consistent with these ATPase motors interacting at the same T4P machine as suggested by the AF3 model.

Our data thus far strongly suggest that PilT and PilU interact with one another to engage the T4P machine, which is consistent with the AF3 model. This is also supported by a recent preprint that analyzes the *V. cholerae* competence T4P machine *in situ* via cryo-EM. In strains expressing *pilT^WA^* and *pilU^WA^*, subtomogram averages revealed the presence of two stacked densities below the T4P machine that nicely fit the PilT-PilU AF3 model (32), which is consistent with stacked PilT and PilU hexamers engaging the T4P machine. Altogether, these results support the ultrastructural organization of the PilC-PilT-PilU complex predicted by the AF3 hybrid model.

### Interactions between PilT and the PilU^C-term^ are critical for T4P retraction

Our data thus far suggest that PilT and PilU interact and that this interaction is likely required for PilU to engage the T4P machine. Next, we sought to employ the PilT-PilU AF3 model to further define the molecular basis for motor ATPase coordination.

Competence T4P retraction is necessary for DNA uptake (1, 11); thus, NT serves as a simple, yet sensitive readout (>5-logs of dynamic range) for T4P retraction (**Fig. 2b**). Competence T4P can also be directly imaged in *V. cholerae* via introduction of a cysteine into the major pilin (*pilA^S67C^*), which allows for subsequent labeling with fluorescent maleimide dyes (11, 33). Strains that are competent for T4P retraction have very few extended pili because competence T4P are highly dynamic (**Fig. 2c**, parent). In contrast, strains that cannot retract competence T4P become hyperpiliated due to the accumulation of static extended pili on the cell surface (**Fig. 2c**, Δ*pilT*). Competence T4P also self-interact (27); thus, hyperpiliated cells form large multi-cellular aggregates that “floc” out of solution (**Fig. 2c**) (18). PilU is normally dispensable for competence T4P retraction, however, if cells contain a PilT^WA^ mutation, retraction becomes dependent on PilU (18, 19), a finding that we recapitulate here (**Fig. 2b-c**, compare Δ*pilU* to *pilT^WA^* Δ*pilU*). Thus, the *pilT^WA^* background can be used to assess PilT-PilU-dependent retraction (*i.e.*, retraction that is dependent on <u>both</u> motor ATPases), while a background where PilT ATPase activity is intact and *pilU* is deleted (*pilT^+^* Δ*pilU*) can be used to assess PilT-dependent retraction (*i.e.*, retraction that is only dependent on PilT). For this reason, we initially focused on mutating the PilT side of the PilT-PilU interface to define regions that are 1) critical for these motors to coordinate and promote PilT-PilU-dependent T4P retraction but 2) dispensable for PilT-dependent retraction. Analysis of the AF3 model revealed three distinct regions of PilT that putatively form salt bridges with PilU – PilT^D2,E53^, PilT^R35,K36^, and PilT^E64,E65^ (**Fig. 2a and Fig. S4**). To test if any of these areas are required for PilT-PilU-dependent retraction, we mutated these PilT residues to alanines and assessed pilus retraction using NT, direct observation of competence T4P, and cellular aggregation as described above. The phenotype of *pilT^WA,E64A,E65A^* was indistinguishable from *pilT^WA^*, suggesting that this region is dispensable for PilT-PilU-dependent retraction (**Fig. 2b-c**). By contrast, *pilT^WA,R35A,K36A^* and *pilT^WA,D2A,E53A^* resulted in an ∼3.5-log decrease in NT (compared to *pilT^WA^*) and a strong hyperpiliation and aggregation phenotype which is consistent with these two regions playing an important role in PilT-PilU-dependent retraction (**Fig. 2b-c**). Importantly, mutating these three regions (*pilT^E64A,E65A^*, *pilT^D2A,E53A^*, and *pilT^R35A,K36A^*) does not diminish PilT- dependent T4P retraction (*i.e.*, retraction in a *pilT*^+^ Δ*pilU* background), indicating that PilT^D2,E53^ and PilT^R35,K36^ are specifically required for PilT-PilU-dependent retraction (**Fig. 2b-c**).

**Fig. 2.**
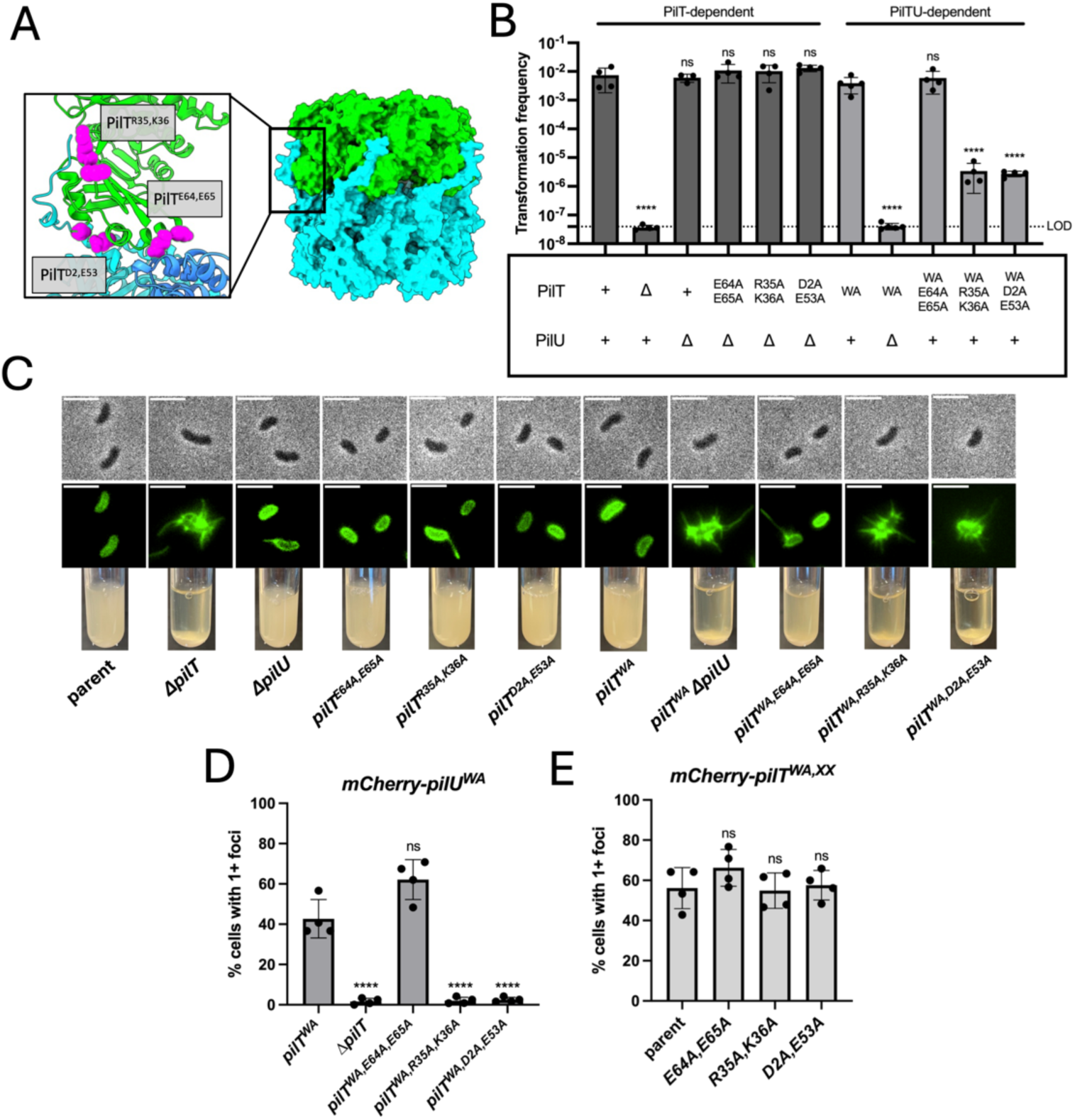
Contacts between PilT and the PilU^C-term^ are critical for PilT-PilU interactions and T4P retraction. (**A**) AF3 model of PilT (green)-PilU (cyan) interactions. Inset highlights the interaction interfaces targeted for mutagenesis. The PilT residues that putatively participate in intermolecular salt bridges are shown in magenta. (**B**) NT assays of the indicated *V. cholerae* strains. Dark gray bars denote strains that report on PilT-dependent retraction (*i.e.*, in a *pilT^+^* background where PilT ATPase activity is intact), while light grey bars denote strains that report on PilT-PilU-dependent retraction (*i.e.*, in a *pilT^WA^* background where retraction is dependent on PilU). LOD, limit of detection. (**C**) Representative images of surface piliation (top panel) and cellular aggregation (bottom panel) for the indicated strains. Scale bars on micrographs, 3 µm. Data are representative of two independent experiments. (**D**) Motor localization assay of cells expressing *mCherry-pilU^WA^*and the indicated *pilT* alleles. For each strain, the percentage of cells (*n* = 120 cells analyzed) with one or more fluorescent mCherry-PilU^WA^ foci is reported. (**E**) Motor localization assay of *ΔpilU* cells expressing an *mCherry-pilT^WA^*construct with the additional mutations indicated. For each strain, the percentage of cells (*n* = 120 cells analyzed) with one or more fluorescent mCherry-PilT^WA^ foci is reported. Data in **B**, **D**, and **E** are from 4 independent biological replicates and shown as the mean ± SD. Statistical comparisons were made by one-way ANOVA with Tukey’s multiple comparison test of the log-transformed data. ns, not significant; **** = *p* < 0.0001. Symbols directly above bars denote comparisons to the appropriate parent strain; in **B**, symbols above the dark grey bars denote comparisons to the parent strain, while symbols above the light grey bars denote comparisons to *pilT^WA^*.

The reduction in PilT-PilU-dependent retraction observed in *pilT^D2A,E53A^* and *pilT^R35A,K36A^* may be due to an effect of these mutations on either (1) diminishing PilT-PilU interactions or (2) preventing PilT-PilU coordination without affecting their interaction. To distinguish between these, we assessed the effect of these mutations on PilT and PilU localization to T4P machines. We hypothesized that if these regions are required for PilT-PilU interactions, mutating these residues in a *pilT^WA^ mCherry-pilU^WA^* background would result in loss of mCherry-PilU^WA^ foci.

When we performed this analysis, *pilT^WA,D2A,E53A^ mCherry-pilU^WA^* and *pilT^WA,R35A,K36A^ mCherry- pilU^WA^* no longer formed mCherry-PilU^WA^ foci (**Fig. 2d**). While *pilT^WA,D2A,E53^* and *pilT^WA,R35A,K36A^* do not completely eliminate PilT-PilU-dependent transformation (**Fig. 2b-c**); transformation is still reduced ∼1000-fold. Thus, while there is likely still some residual PilT-PilU interactions in these backgrounds, it is below the limit of detection of our microscopy-based motor localization assay which inherently has a narrower dynamic range. By contrast, *pilT^WA,E64A,E65A^ mCherry-pilU^WA^* still formed mCherry-PilU^WA^ foci and was indistinguishable from the *pilT^WA^ mCherry-pilU^WA^* parent (**Fig. 2d**), which is consistent with this mutation having no effect on PilT-PilU-dependent retraction (**Fig. 2b-c**). Importantly, mCherry-PilT^WA,E64A,E65A^, mCherry-PilT^WA,D2A,E53A^, and mCherry- PilT^WA,R35A,K36A^ all localized to T4P machines just like the mCherry-PilT^WA^ parent (**Fig. 2e**), which is consistent with these mutations having no effect on PilT-dependent retraction (**Fig. 2b-c**). Together, these data suggest that PilT^D2,E53^ and PilT^R35,K36^ are important for the interaction between PilT and PilU that allow this complex to coordinate retraction.

PilT^D2,E53^ and PilT^R35,K36^ are both predicted to interact with the PilU C-terminus (PilU^C-term^). PilT and PilU are highly homologous motor ATPases, however the extended C-terminal domain is a unique feature of PilU homologs (**Fig. S5**). The AF3 model predicts that the PilU^C-term^ wraps around the sides of the PilT hexamer (**Fig. 2a**). Because our mutational analysis demonstrates that PilT^D2,E53^ and PilT^R35,K36^ diminish PilT-PilU-dependent retraction (**Fig. 2**), we hypothesized that interactions between PilT and the PilU^C-term^ are critical for PilT-PilU mediated T4P retraction.

To test this hypothesis at a higher resolution, we targeted the putative intermolecular salt bridges formed between PilT^D2,E53^-PilU^K348^, PilT^R35-^PilU^D366,E368^, PilT^K36^-PilU^D366^ for mutagenesis by flipping their charge (**Fig. 3a**). We reasoned that flipping the charge on either side of the intermolecular salt bridge should have a repulsive effect that breaks PilT-PilU interactions, which should disrupt pilus retraction. However, flipping the charge on both sides of the salt bridge should repair the interaction and restore T4P retraction. Importantly, restoration of T4P retraction in the double charge-flip mutant should only occur if the two residues interact, thus, providing an easy genetic approach for testing the residue-level accuracy of the PilT-PilU interactions predicted by AF3.

**Fig. 3.**
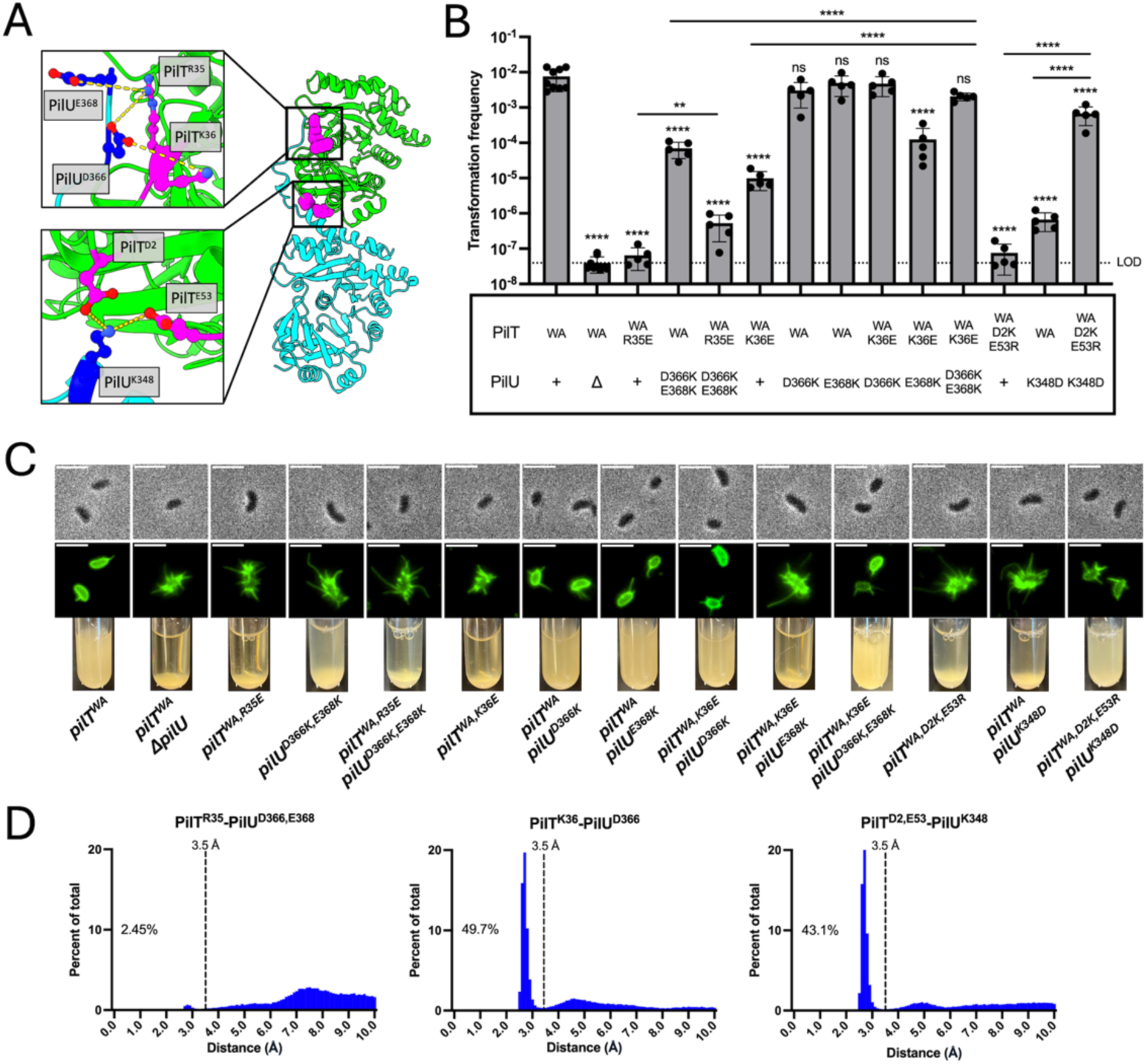
AF3-MD simulations and high-resolution genetic analysis reveal PilT-PilU^C-term^ salt bridges required for motor coordination. (**A**) AF3 model of PilT (green)-PilU (cyan) interactions. The inset highlights the residues that participate in the three putative salt bridges targeted for mutagenesis. PilT residues are shown in magenta with nitrogen atoms colored blue, and PilU residues are shown in dark blue with oxygen atoms colored red. (**B**) NT assay of the indicated *V. cholerae* strains. Data are from five independent biological replicates and shown as the mean ± SD. LOD, limit of detection. Statistical comparisons were made by one-way ANOVA with Tukey’s multiple comparison test of the log-transformed data. ns, not significant; ** = *p* < 0.01; **** = *p* < 0.0001. Symbols directly above bars denote comparisons to the *pilT^WA^* parent. (**C**) Representative images of surface piliation (top panel) and cellular aggregation (bottom panel) for the indicated strains. Scale bar for micrographs, 3 µm. Data are representative of two independent experiments. (**D**) Distance frequency histogram for the indicated residue pairs during the MD simulation of the AF3 PilT-PilU model. Distances between the side chains of the indicated residues were measured at 0.1-ns intervals over the 500-ns simulation. Percentage on graph denotes the amount of time that the indicated side chains are <3.5 Å (dotted line). Data are compiled from two independent MD simulations.

As shown above, mutating the PilT side of the interface allows us to test whether mutations specifically affect PilT-PilU-dependent retraction (tested in the *pilT^WA^* background) without disrupting PilT-dependent retraction (tested in the *pilT^+^* Δ*pilU* background). By contrast, mutating the PilU side of the interface cannot distinguish between 1) disruption of the PilT-PilU interface and 2) mutations that decrease PilU stability or cause it to misfold. Thus, we first assessed the impact of flipping the charge on the PilT side of the interface in the three predicted salt bridge contacts (*pilT^R35E^*, *pilT^K36E^*, and *pilT^D2K^*^,*E53R*^). All three mutants resulted in a decrease in NT, with *pilT^R35E^* and *pilT^D2K^*^,*E53R*^ causing the most severe loss of retraction (**Fig. 3b-c**). Importantly, these mutations had no observable impact on PilT-dependent retraction (**Fig. S6**). Thus, disrupting the PilT side of the putative PilT^D2,E53^-PilU^K348^, PilT^R35^-PilU^D366,E368^, and PilT^K36^- PilU^D366^ salt bridges specifically inhibited PilT-PilU-dependent T4P retraction.

To further probe the validity of these intermolecular salt bridges, we sought to flip the charge on the PilU side of the interface to test whether this recovered T4P retraction. The AF3 model indicates that PilT^R35^ makes contacts with both PilU^D366^ and PilU^E368^ (**Fig. 3a**). The *pilT^WA,R35E^ pilU^D366K,E368K^* strain, however, remained hyperpiliated and only had a modest ∼1 log restoration in NT when compared to *pilT^WA,R35E^ pilU^WT^* (**Fig. 3b-c**). Importantly, *pilU^D366K,E368K^* did not affect steady-state levels of PilU (**Fig S7a**). Together, these results suggest that PilT^R35^ does not form a salt bridge with PilU^D366,E368^. To more rigorously examine the stability of these putative intermolecular salt bridges, we performed all-atom molecular dynamics (MD) simulations of the PilT-PilU AF3 model. The resulting 500-ns MD simulations were then analyzed to determine the percent of time that residues remained in close enough proximity to form a salt bridge (<3.5 Å), with a higher percentage indicating a more stable interaction. The AF3-MD simulations indicated that PilT^R35^ was not predicted to form a stable interaction with either PilU^D366^ or PilU^E368^ (**Fig. 3d**, **S8,** and **Movie S10**). Thus, while PilT^R35^ is clearly critical for PilT-PilU-dependent retraction, it likely does not form a salt bridge with PilU. Instead, PilT^R35^ may promote PilT-PilU interactions via other intermolecular contacts. Accordingly, the MD simulation suggests that PilT^R35^ interacts with PilU^I365^ and PilU^M367^ (**Fig. S9** and **Movie S11**).

For PilT^K36^, the AF3-MD simulation predicted a stable salt bridge for PilT^K36^-PilU^D366^ (**Fig. 3d, S8,** and **Movie S12**). Consistent with this, introducing *pilU^D366K^* into the *pilT^WA,K36E^* background restored T4P retraction to parent levels (**Fig. 3b-c**). Importantly, steady-state levels of PilU^D366K^ are similar to the PilU^WT^ parent (**Fig S7a**). If PilT^K36^ and PilU^D366^ exclusively form a salt bridge, a charge flip mutation to either residue should result in a similar transformation deficit. Unlike *pilT^WA,K36E^*, however, *pilT^WA^ pilU^D366K^* transforms like the parent (**Fig. 3b-c**). This suggests that PilT^K36^ may form additional contacts with PilU. The AF3-MD simulations suggest that PilT^K36^ may interact with PilU^E368^; and these interactions were not altered when we performed MD simulations of the PilT-PilU^D366K^ complex (**Fig. S10**). Based on this, we hypothesized that PilT^K36^- PilU^E368^ may partially compensate for the loss of PilT^K36^-PilU^D366^ interactions in the PilU^D366K^ background. Consistent with this, *pilT^WA^ pilU^D366K,E368K^* exhibited a ∼2-log transformation deficit compared to *pilT^WA^ pilU^D366K^* (**Fig. 3b**). If this difference is due to the loss of residual PilT^K36^- PilU^E368^ interactions, then we hypothesized that the transformation deficit of *pilT^WA^ pilU^D366K,E368K^* should be completely restored to parent levels in *pilT^WA,K36E^ pilU^D366K,E368K^* (similar to what is observed for *pilT^WA,K36E^ pilU^D366K^*). Indeed, *pilT^WA,K36E^ pilU^D366K,E368K^* exhibited a transformation phenotype that was indistinguishable from the parent (**Fig. 3b**). Thus, PilT^K36^ likely interacts with both PilU^D366^ and PilU^E368^; with PilT^K36^-PilU^D366^ interactions playing the dominant role since recovery of transformation is greater in *pilT^WA,K36E^ pilU^D366K^* than in *pilT^WA,K36E^ pilU^E368K^* (**Fig. 3b**).

For PilT^D2,E53^, the AF3-MD simulation predicted a stable salt bridge between PilT^D2,E53^-PilU^K348^, with all three residues participating in the interaction (**Fig. 3d, S8,** and **Movie S13**). Accordingly, *pilT^WA,D2K,E35R^ pilU^K348D^* largely restored NT and surface piliation when compared to *pilT^WA,D2K,E35R^ pilU^WT^ and pilT^WA^ pilU^K348D^* (**Fig. 3b-c**). Importantly, steady-state levels of PilU^K348D^ are similar to the PilU^WT^ parent (**Fig. S7a**).

Together, these results help further validate the AF3 model and demonstrate that PilT-PilU^C-term^ interactions are critical for PilT-PilU-dependent retraction at the molecular scale.

### PilT-PilU^C-term^ contacts are necessary for the high retraction forces needed for NT under physiologically relevant conditions

Thus far, we have used the *pilT^WA^* background to isolate and study PilT-PilU-dependent retraction, but when PilT ATPase activity is intact, PilU is dispensable for competence T4P retraction. Accordingly, Δ*pilU* can take up soluble transforming DNA (tDNA) from the environment at a similar frequency to the parent (**Fig. 2b** and **4a**). PilU, however, is critical for generating high retraction forces (18, 20). We recently demonstrated that PilU-dependent retraction is necessary for NT when tDNA is adsorbed to a chitin surface (24), a finding that we recapitulate here (**Fig. 4b-c**). Thus, assessing NT of *V. cholerae* in chitin biofilms under flow allows us to study PilT-PilU-dependent retraction in a physiologically relevant setting.

**Fig. 4.**
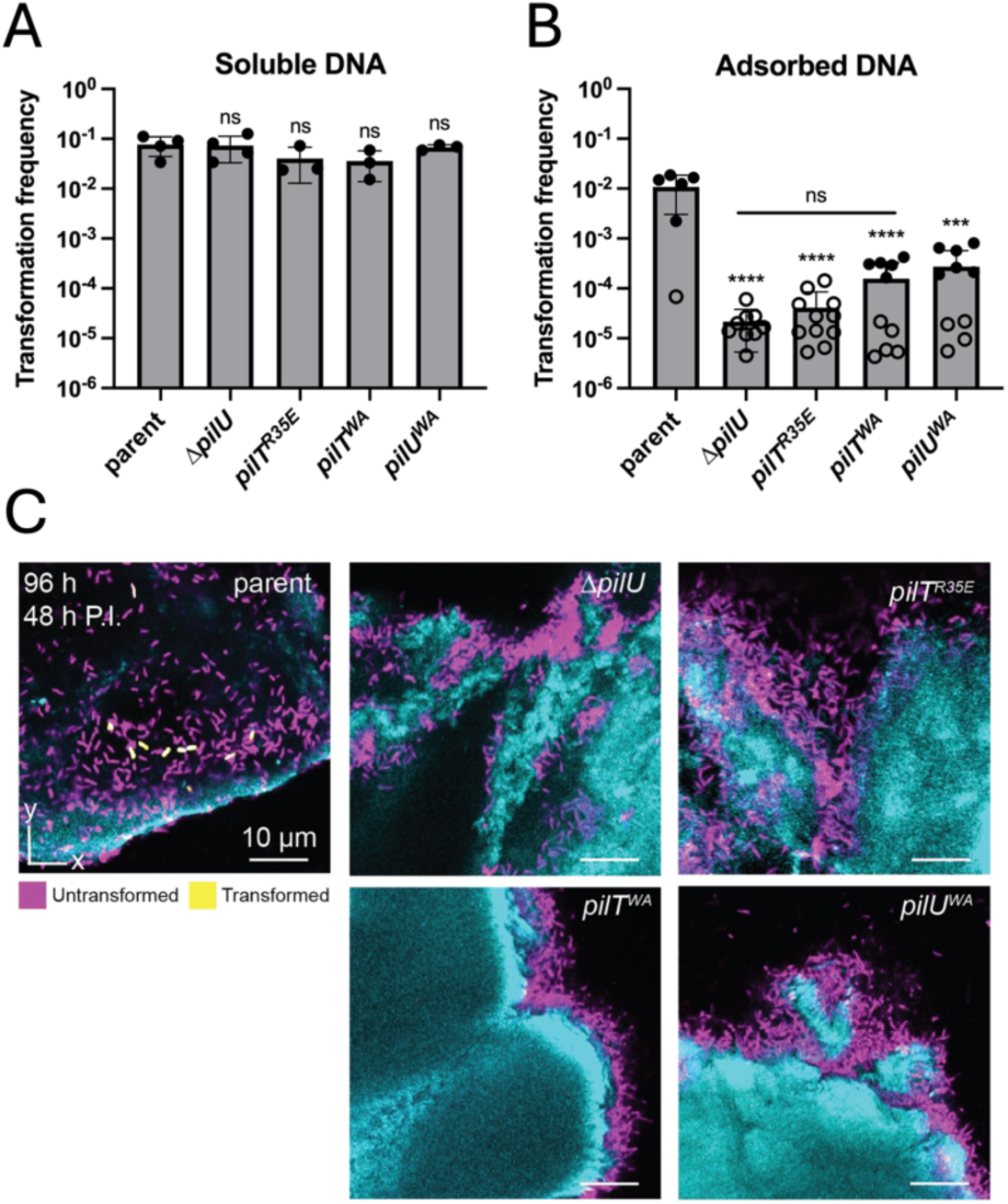
PilT-PilU^C-term^ interactions are critical for motor coordination under physiologically relevant conditions. (**A**) Chitin-dependent NT assays of the indicated *V. cholerae* strains with soluble tDNA. (**B**) Flow biofilm NT assays where tDNA was adsorbed to the chitin surface. Open circles denote replicates where no transformants were observed and the limit of detection for that sample is plotted. (**C**) Representative images of assays described in **B** highlighting transformed (yellow) and untransformed (magenta) cells within the chitin-bound biofilm. The chitin surface is shown in cyan and appears irregular in shape due to optical sectioning through the 3-dimensional shrimp shell particles used for these experiments. All data are from at least 3 independent biological replicates and shown as the mean ± SD. Statistical comparisons were made by one-way ANOVA with Tukey’s multiple comparison test of the log-transformed data. ns, not significant; *** = *p* < 0.001; **** = *p* < 0.0001. Symbols directly above bars denote comparison to the parent.

Based on the results described above, we hypothesized that PilT-PilU^C-term^ contacts would be critical for coordinating the forceful T4P retraction required for NT in chitin flow biofilms when tDNA is adsorbed to chitin. Consistent with this, *pilT^R35E^*, the single charge-flip mutation that had the strongest impact on PilT-PilU-dependent retraction (**Fig. 3b**), exhibited a loss in NT that was indistinguishable from Δ*pilU* when tDNA was adsorbed to chitin (**Fig. 4b-c**). Importantly, the ATPase activity of both PilT and PilU are intact in this mutant; thus, the disruption of PilT-PilU^C- term^ interactions in *pilT^R35E^* likely disrupts the ability of PilT-PilU to generate the high retraction forces required for uptake of chitin-adsorbed tDNA. Consistent with this, *pilT^R35E^* did not impact NT when tDNA was freely soluble (**Fig. 4a**), suggesting that PilT-PilU^C-term^ contacts are specifically required for forceful T4P retraction. Together, these results suggest that PilT-PilU^C-term^ contacts are critical for these motors to coordinate under physiologically relevant conditions where forceful T4P retraction is necessary for DNA uptake.

While these data suggest that PilT-PilU^C-term^ contacts are important for high force retraction, it remains unclear why coordination between PilT and PilU is necessary to generate high retraction forces. Our results thus far indicate that PilT-PilU form stacked hexamers that are arranged head-to-tail, and that PilU^C-term^ binds to the sides of the PilT hexamer in a manner that could allow PilU to “pull” on PilT during its catalytic cycle. It is possible this arrangement allows these motors to intimately coordinate their catalytic cycles to combine their efforts and generate forces that could not be achieved by either motor alone. To explore this idea, we sought to estimate the maximum force that a single T4P retraction ATPase can generate (see **Methods** for details). This estimate is derived from four main assumptions: 1) that the free energy of ATP hydrolysis under physiological conditions is -47 kJ/mol (34), 2) that two molecules of ATP are hydrolyzed per catalytic cycle (12, 35), 3) that each catalytic cycle removes one pilin subunit from the pilus filament resulting in a step size of 1 nm (13, 36), and 4) that AAA+ motor ATPases like PilT/PilU operate at an efficiency of ∼32.5% (37, 38). Based on this, we estimate that a single T4P retraction motor can generate ∼50 pN of force. T4P, however, have empirically been shown to retract at forces that exceed 100 pN (39–41). Thus, molecular coordination between PilT-PilU may, in fact, be necessary to achieve the high T4P retraction forces observed in nature.

According to this model, the ATPase activity of both PilT and PilU should be necessary to generate high retraction forces. To test this, we assessed the transformation of strains where the ATPase activity of either PilT or PilU is ablated via a Walker A mutation (*pilU^WA^* and *pilT^WA^*) in chitin flow biofilms where high force retraction is required for DNA uptake. Consistent with our model, both *pilT^WA^* and *pilU^WA^* had reduced transformation frequencies compared to the parent when the tDNA was adsorbed to the chitin (**Fig. 4b-c**). Importantly, transformation was indistinguishable from the parent when the tDNA was freely soluble (**Fig. 4a**). Together, these data suggest that the ATPase activity of both PilT and PilU is required for maximal T4P retraction force. Interestingly, *pilU^WA^* did not phenocopy the transformation deficit of Δ*pilU* in these assays (**Fig. 4b-c**). This may suggest that PilU can also promote forceful retraction in a manner that is independent of its ATPase activity (see **Discussion**).

### PilT-PilU^C-term^ contacts are important for T4P retraction in Acinetobacter baylyi

PilU is broadly conserved among gammaproteobacteria including, but not limited to, *Acinetobacter baylyi*, *Pseudomonas aeruginosa*, *Legionella pneumophila*, and *Xylella fastidiosa* (**Fig. S5a**). Thus, we next sought to determine whether PilT-PilU^C-term^ contacts are a conserved mechanism for PilT-PilU coordination in diverse T4P systems. Sequence alignments highlight that while the majority of PilU^C-term^ is not conserved, the PilU_Vc_^K348^ residue that forms a salt bridge with PilT_Vc_^D2,E53^ is conserved in all PilU homologs (**Fig. S5a**). Accordingly, the PilT_Vc_^D2,E53^- PilU_Vc_^K348^ salt bridge is conserved in AF3 predictions of these PilT-PilU homologs (**Fig. S5b**).

*Acinetobacter baylyi* competence T4P represent another well-established model system for studying T4P (18, 25, 42–45). For this T4P system, NT is also a simple and sensitive readout (∼4- logs of dynamic range) for pilus retraction (**Fig. 5b**). In addition, prior work demonstrates that *pilU_Ab_* is critical for T4P retraction in a *pilT_Ab_^WA^* background (18), a finding that we recapitulate here (**Fig. 5b**).

**Fig. 5.**
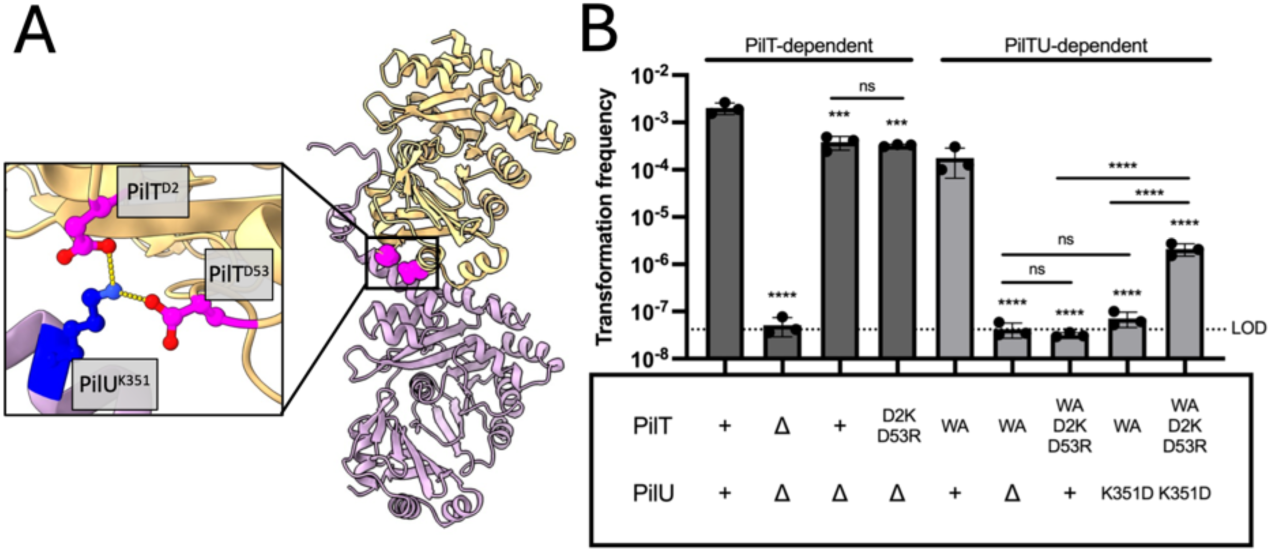
PilT-PilU^C-term^ contacts are important for T4P retraction in *A. baylyi*. (**A**) AF3 model of PilT_Ab_ (orange)- PilU_Ab_ (pink) interactions. Inset shows the conserved PilT_Ab_^D2,D53^-PilU_Ab_^K351^ salt bridge. (**B**) NT assays of the indicated *A. baylyi* strains. Dark gray bars denote strains that report on PilT-dependent retraction (*i.e.*, in a *pilT^+^* background where PilT ATPase activity is intact), while light grey bars denote strains that report on PilT-PilU-dependent retraction (*i.e.*, in a *pilT^WA^* background where retraction is dependent on PilU). Data are from three independent biological replicates and shown as the mean ± SD. Statistical comparisons were made by one-way ANOVA with Tukey’s multiple comparison test of the log-transformed data. ns, not significant; *** = *p* < 0.001; **** = *p* < 0.0001. Symbols above the dark grey bars denote comparisons to the parent, while symbols above the light grey bars denote comparisons to *pilT^WA^*.

The PilT_Ab_^D2,D53^-PilU_Ab_^K351^ contact in *A. baylyi* (**Fig. 5a**) is equivalent to the PilT_Vc_^D2,E53^-PilU_Vc_^K348^ salt bridge that facilitates PilT-PilU-dependent retraction in *V. cholerae* (**Fig. S5a-b**). To test whether PilT-PilU^C-term^ contacts are also required for the coordination of these motor ATPases in *A. baylyi*, we targeted the PilT_Ab_^D2,D53^-PilU_Ab_^K351^ interface for mutational analysis. Flipping the charge on either side of the interface (*pilT^WA,D2K,D53R^ pilU^WT^* or *pilT^WA^ pilU^K351D^*) results in an NT deficit that is indistinguishable from *pilT^WA^* Δ*pilU* (**Fig. 5b**). Importantly, *pilT^D2K,D53R^* did not impact PilT-dependent retraction (**Fig. 5b**; compare *pilT^D2K,D53R^* Δ*pilU* to *ΔpilU*), and steady-state levels of PilU^K351D^ were similar to the PilU^WT^ parent (**Fig. S7b)**. Flipping the charge on both sides of the interface (*pilT^WA,D2K,^*^D53R^ *pilU^K351D^*) partially (∼1.5-log) restores NT (**Fig. 5b**), which strongly suggests that these residues form a salt bridge. Together, these results indicate that PilT-PilU^C- term^ contacts are a conserved mechanism for PilT-PilU coordination in diverse T4P.

## DISCUSSION

While prior work established that PilU requires PilT to facilitate retraction (18, 19), a molecular explanation for this interdependence was lacking. Here, we show that PilT is required to “bridge” PilU interactions with the T4P machine *in vivo*. By contrast, PilT could engage T4P machines even in the absence of PilU. PilT homologs contain a highly conserved “AIRNLIRE” motif at the C-terminus (**Fig. S5a**) (46). The PilT-PilC AF3 model places the AIRNLIRE motif within the PilT-PilC interface (**Fig. S11**)(47), suggesting that it may be required for PilT-PilC interactions. Interestingly, the AIRNLIRE motif is not conserved among PilU homologs (**Fig. S5**), which may be one reason why PilU does not directly engage the PilC platform. Our results also provide insight into *how* PilT and PilU interact to coordinate retraction. Specifically, we show that interactions between PilT and the PilU^C-term^, a feature that is unique to PilU homologs (**Fig. S5a**), is critical for these proteins to promote T4P retraction. The strong conservation between PilT and PilU indicates that PilU likely arose through duplication of PilT and subsequent evolution of the resulting paralog (12, 48). Thus, it is tempting to speculate that the emergence of PilT-PilU coordination required loss of the “AIRNLIRE” motif and gain of the C-terminal extension in PilU.

While our results show that PilT-PilU^C-term^ interactions are critical for motor coordination, it is important to acknowledge that this is likely only one of the many molecular interactions that are required for these motors to coordinate. Prior work demonstrates that T4P motor ATPases likely adopt a C2 symmetry and undergo dynamic conformational changes throughout their catalytic cycle (12, 35, 49). AF3, however, only models PilT and PilU with C6 symmetry, highlighting that these models cannot provide insight into the molecular interactions required for motor coordination throughout the PilT-PilU catalytic cycle. Future work will focus on integrating the catalytic dynamics of these motor proteins with the PilT-PilU^C-term^ interactions identified in our study.

Although our results provide insight into the molecular coordination between PilT-PilU, it remains unclear why this coordination is necessary. Here, we propose a model where coordination of the catalytic cycles of both PilT and PilU is necessary to generate the high T4P retraction forces observed in nature. Consistent with this, we show that the ATPase activity of both PilT and PilU contributes to tDNA uptake in conditions that require forceful retraction. Interestingly, our results suggest that PilU may promote forceful T4P retraction in multiple ways: 1) in an ATPase-dependent manner (**Fig. 4b**; compare parent to *pilU^WA^*) and 2) in an ATPase-independent manner (**Fig. 4b**; compare *pilU^WA^* to Δ*pilU*). One explanation for this ATPase-independent effect is that PilU allosterically stimulates PilT activity (*e.g.*, by increasing the mechanical work output of PilT’s ATPase cycle to enhance its efficiency). This hypothesis, however, is challenging to test empirically because purified PilT and PilU have no detectable ATPase activity *in vitro* (18), which may suggest that motor-T4P machine interactions are required for ATPase activity.

Motor ATPase coordination is a prevalent feature among diverse T4P systems. Several gammaproteobacteria encode PilT and PilU homologs (**Fig. S5**), and we demonstrate that the broadly conserved PilT^D2,E53^-PilU^K348^ salt bridge is important for PilT-PilU coordination in both *V. cholerae* and *A. baylyi*. While PilT^R35^ and PilT^K36^ are also conserved among PilT homologs (**Fig. S5a**), the specific contacts between these residues and PilU are not conserved; in fact, the C-terminal region of PilU past K348 is poorly conserved (**Fig. S5a**). This suggests that PilT-PilU coordination may be mediated by both conserved and species-specific PilT-PilU^C-term^ interactions. Consistent with this, PilU homologs from different species cross-complement with varying degrees of efficacy (26). Outside of the gammaproteobacteria, some species encode multiple PilT paralogs (*e.g.*, *Neisseria spp.* and *Myxococcus spp.*), suggesting that motor ATPase coordination may also be conserved in more distantly related T4P systems. Additionally, some T4P rely on multiple motor ATPases for pilus extension (42, 50), suggesting that motor coordination may be a critical feature of both T4P extension and retraction.

## METHODS

### Bacterial strains and culture conditions

*Vibrio cholerae* and *Acinetobacter baylyi* strains were routinely grown in LB broth and on LB agar plates. Medium was supplemented with kanamycin (50 µg/mL), spectinomycin (200 µg/mL), sulfamethoxazole (100 µg/mL), trimethoprim (10 µg/mL), carbenicillin (100 µg/mL), and chloramphenicol (1 µg/mL) when appropriate. A comprehensive list of strains used in this study can be found in **Table S1**.

### Natural Transformation (NT) assays

Chitin-independent NT assays of *V. cholerae* were conducted using strains where 1) TfoX, the master competence regulator, was placed under an IPTG-inducible promoter (P_tac_-*tfoX*), and 2) *luxO* was deleted to genetically lock cells in a high-density state. To induce natural competence, strains were grown to late-log by rolling at 30 °C in LB supplemented with 20 mM MgCl_2_, 10 mM CaCl_2_, and 100 µM IPTG. Then, 7 µL of this culture was diluted into 350 *µL* of Instant Ocean medium (IO; 7 g/L Aquarium Systems) for each transformation reaction. Two reactions were setup for each strain: one where 100 ng of tDNA (a ΔVC1807::Kan^R^ PCR product) was added and a negative control where no tDNA was added. After 3 min of tDNA incubation, 6 units of DNase I (NEB) was added to stop DNA uptake, and reactions were incubated overnight at 30 °C to allow for tDNA integration. The next day, 0.5 mL LB was added, and reactions were outgrown at 37 °C shaking for 3 hrs. Reactions were then dribble plated for quantitative culture on LB plates with kanamycin (to quantify transformant CFUs) and on plain LB agar plates (to quantify total CFUs). The transformation frequency was determined by dividing the CFUs of transformants by the total CFUs. For reactions with no transformants, the limit of detection was calculated and plotted.

Chitin-dependent NT assays of *V. cholerae* in the presence of soluble DNA or chitin adsorbed DNA were conducted exactly as previously described (24). Briefly, strains contained two fluorescent reporters: a disrupted GFP construct (ΔVC1807::Cm^R^-*lacI*-P_tac_*-gfp::parST1*) and a constitutively expressed mKate2 construct (Δ*lacZ*::Spec^R^-*lacI*-P_tac_-*mKate2*).

For assays where free DNA was supplied, strains were grown rolling at 30 °C in LB to mid-log. Cells were then washed and adjusted to an OD_600_ = 1.0 in IO. Next, chitin-dependent transformation reactions were setup by mixing the following in a 2mL eppi: 100 µL of cells, 150 µL chitin slurry (8 g / 150 mL IO; Alfa Aesar), and 750 µL IO. Two reactions were prepped for each strain. Reactions were incubated static overnight at 30 °C to induce natural competence. The next day, 1000 ng of tDNA (a ΔVC1807::Cm^R^-*lacI*-P_tac_*-gfp* PCR product) was added to one reaction, while no DNA was added to the other reaction, which served as a negative control. All reactions were incubated overnight at 30 °C with shaking to ensure the tDNA remained in solution. The next day, cells were harvested from the chitin, placed under a 0.4% IO gelzan pad and imaged via epifluorescence microscopy with FITC and mCherry filter cubes. Images were analyzed using MicrobeJ to determine the number of mKate (total) and GFP (transformant) fluorescent cells. Transformation frequency was calculated by dividing the GFP (transformant) cell counts by the mKate (total) cell counts (at least 500 cells analyzed in each sample).

For assays where tDNA was chitin adsorbed, flow cells (24) containing chitin flakes were first washed with 0.13 ng/μL of tDNA (a ΔVC1807::Cm^R^-*lacI*-P_tac_*-gfp* PCR product) at a volumetric flow rate of 0.1 μL/min for 24 h. Following this tDNA adsorption step, the flow cell inlet tubing was swapped to blank defined artificial seawater (DASW; consists of 234 mM NaCl, 27.5 mM MgSO4, 1.5 mM NaHCO3, 4.95 mM CaCl2, 5.15 mM KCI, 0.07 mM Na2B4O7, 0.05 mM SrCl, 0.015 mM NaBr, 0.001 mM NaI, 0.013 mM LiCl, 0.187 mM K2HPO4, and pH 7.1 triethanolamine) and flow was maintained at 0.1 μL/min for an additional 24 h to remove any unbound tDNA. After this wash step, 100 μL of reporter cells were resuspended in DASW to an OD_600_ of 5.0 and inoculated into the chambers via syringe, after which flow with blank DASW was resumed. The reporter strain was then incubated at room temperature (∼22 °C) for 48 h before imaging. Flow cells were imaged using a Zeiss 880 line-scanning confocal microscope and a 40x/1.2 N.A. water objective. A 488 nm laser line was used to excite the GFP protein expressed after transformation events and a 594 nm laser line was used to excite the mKate2 protein carried by the reporter strain. BiofilmQ was used to threshold images and quantify the reporter (mKate2 labeled) and transformant (GFP labeled) biovolumes (51). The transformation frequency was calculated as GFP biovolume divided by mKate2 biovolume. For samples where no transformants were observed, the approximate biovolume of a single cell was used to calculate the limit of detection and plotted.

For NT assays of *A. baylyi*, overnight cultures were washed and adjusted to an OD_600_ = 2.0 in fresh LB. Then, 50 µL of this culture was added to 450 µL LB supplemented with 20 mM MgCl_2_ and 10 mM CaCl_2_ in a 2 mL eppi. Two transformation reactions were setup for each strain: one where 50 ng of tDNA (a ΔACIAD1551::Kan^R^ PCR product) was added and a negative control where no tDNA was added. Transformations were incubated with end-over-end rotation at 30°C for 45 mins. After 45 min of tDNA incubation, 6 units of DNase I (NEB) was added to stop DNA uptake, and reactions were incubated with end-over-end rotation at 30 °C for an additional 5 hrs to allow for tDNA integration and outgrowth. Reactions were then dribble plated for quantitative culture on LB plates with kanamycin (to quantify transformant CFUs) and on plain LB agar plates (to quantify total CFUs). The transformation frequency was determined by dividing the CFUs of transformants by the total CFUs. For reactions with no transformants, the limit of detection was calculated and plotted.

### Construction of mutant strains

All *V. cholerae* strains used in this study were derivatives of the El Tor isolate E7946 (52). All *A. baylyi* strains were derivatives of strain ADP1 (53). All mutant strains were generated by NT (as described above). Mutant constructs were generated by splicing-by-overlap-extension (SOE) exactly as previously described (54–56). A list of all primers used to generate mutant constructs can be found in **Table S2**.

### Microscopy

Unless otherwise indicated, phase and epifluorescence (FITC, YFP, mCherry channels) images were collected on an inverted Nikon Ti-2 microscope with a Plan Apo ×60 objective fitted with a Hamamatsu ORCAFlash 4.0 camera using Nikon NIS Elements imaging software.

### Motor localization assay

For these assays, all *V. cholerae* strains contained the P_tac_-*tfoX* and Δ*luxO* mutations required for chitin-independent induction of competence as describe above for NT assays. Cells were grown overnight in LB supplemented with 20 mM MgCl_2_, 10 mM CaCl_2_, and 100 µM IPTG rolling at 30°C. The next day, 500 µL of overnight culture was subcultured into 3 mL of LB supplemented with 20 mM MgCl_2_, 10 mM CaCl_2_, and 100 µM IPTG and grown for 1 hr rolling at 30 °C. Then, 100 µL of this culture was spun down and washed in IO. Cells were then placed under a 0.4% IO gelzan pad and imaged via timelapse microscopy (2 sec intervals for 10 sec duration) with the phase contrast and mCherry channels. Quantification of fluorescent foci was performed using MicrobeJ features detection. Foci were only counted if they moved less than 0.3 µm between frames and were present in all frames of the timelapse.

For quantification of colocalizing foci, cells were imaged via timelapse microscopy (2 sec intervals for 10 sec duration) with the phase contrast, FITC, and mCherry channels. For colocalizing PilT and PilU foci, MicrobeJ features detection was used to identify cells with GFP- PilU foci. Then, the presence of a colocalizing PilT-mCherry focus was determined manually. The results are presented as the percentage of GFP-PilU foci that colocalized with mCherry-PilT foci. For motor and PilQ foci, the MicrobeJ colocalization feature was used to identify colocalizing motor and msfGFP-PilQ foci that were less than 0.4 µm apart. Results are presented as the percentage of motor foci that colocalized with msfGFP-PilQ foci.

### Pilus labeling and imaging

*V. cholerae* strains were grown exactly as described above for chitin-independent NT assays. Then, 100 µL of late-log cells were pelleted at 8000 x g for 1 min and washed in IO. Cells were then labeled with 25 µg/mL AF488-mal for 15 mins static at room temperature. Cells were then washed three times in IO, ensuring to spin at 8000 x g for 1 min to prevent shearing of surface pili. Cells were then placed under a 0.4% IO gelzan pad and imaged with the phase and FITC channels. Representative images of the piliation phenotype for each strain were selected from the raw microscopy images.

### T4P-dependent aggregation assay

*V. cholerae* strains were grown exactly as described above for chitin-independent NT assays. Subcultures were allowed to sit static at room temperature on the benchtop for 15 mins to let cellular aggregates settle out of solution. Representative images were taken of each strain without disturbing settled cells.

### AF3-MD analysis

The initial structure of the PilT-PilU complex was generated using AlphaFold 3 (28). Propka3 was used to assign the charges of the ionizable residues and the protonation state of histidine residues (57). PilU^D253^ and PilU^E280^ were protonated. Based on their local electrostatic environment, PilT^H151^, PilT^H166^, PilT^H183^, PilT^H229^, PilT^H280^, PilU^H164^, PilU^H240^, and PilU^H276^ were set to HSE (histidine-epsilon). All other histidine residues were set to HSD (histidine-delta).

The PilT-PilU structure was subsequently solvated, using CHARMM-GUI, v3.7 generating a 162Å^3^ water box (58, 59). Potassium (K^+^) and chloride (Cl^−^) ions were added in order to neutralize the system as well as produce an ionic strength of 150 mM. Simulations were conducted with NAMD3 (60) using the CHARMM36m protein force field (61), and TIP3P water model (62). The initial structure was minimized for 10,000 steps, followed by a two-stage equilibration: first, 1ns of MD was run while restraining the protein heavy atoms, and second, 4ns of MD was run while restraining only the protein backbones, thereby equilibrating the box dimensions. Production runs were performed using hydrogen mass repartitioning (HMR), allowing for a uniform 4 fs time step (63, 64).

The simulations were run at a constant temperature (310 K) and constant isotropic pressure (1 atm), maintained by a Langevin thermostat and piston, respectively (65). Long-range electrostatics were calculated at every time step using the particle-mesh Ewald method (66). We set a short-range cutoff for Lennard-Jones interactions at 12 Å with a switching function starting at 10 Å. Two independent replicas were performed for 500 ns.

Animations were rendered with VMD v1.9.4a51 (67), and the distance calculations were performed with MDAnalysis (68, 69), where the minimum distance between two residue side chains was taken as the closest pair of heavy atoms. In the case where multiple interactions are considered (e.g. PilT^R35^-PilU^D366,E368^), the minimum distance was taken for either residue in the target pair. Salt bridges and contacts were defined when this distance was less than a cutoff of 3.5 Å or 4.5 Å, respectively.

Simulations of the PilT-PilU^D366K^ AF3 complex were performed as described above except the number of positive ions was adjusted to ensure neutrality of the simulation cell.

### Estimating T4P force generation

The following equation was used to calculate the maximal force (*F*_*max*_) generated by a T4P motor ATPase:

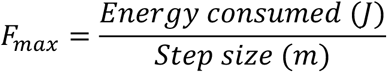

The energy consumed was determined based on two assumptions: 1) the free energy of ATP hydrolysis in physiological conditions is -47 kJ/mol (34) and 2) that two molecules of ATP are hydrolyzed per ATPase cycle (12, 35). Based on this, the free energy released upon hydrolysis of two ATP molecules was determined to be 1.56 x 10^-19^ J. To estimate the step size, we assumed that one pilin is removed from the pilus filament in each ATPase cycle resulting in a step size of 1 nm (or 1 x 10^-9^ m) (13, 36). These values were then plugged into the equation above to calculate the *F*_*max*_ generated by T4P motors at 100% efficiency:

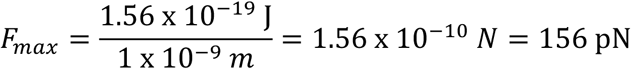

It is unlikely that these motor proteins operate at 100% efficiency. Prior work has defined the efficiencies of two other members of the AAA+ family of motor ATPases: the phi29 packing motor = 30% (37) and ClpX = 35% (38). We averaged these values (32.5%) to generate an estimate for the efficiency of this family of motor ATPases. Accounting for average efficiency, we estimated that a single T4P motor can maximally generate ∼50.7 pN of force.

### Generating multiple sequence alignments

Multiple sequence alignments (MSA) were generated by first gathering the PilT and PilU protein sequences from each organism from GenBank. Sequences were then aligned using T-Coffee (70), and the alignments were reformatted for the final figure using the Sequence Manipulation Suite: Color Align Conservation tool (71).

### Western bloĖng

Strains of *V. cholerae* and *A. baylyi* were grown exactly as described above for natural transformation assays for each respective organism. Cultures were then washed and resuspended to an OD_600_ = 100 in IO. Cells were mixed 1:1 with 2x SDS-PAGE sample buffer (200 mM Tris pH 6.8, 25% glycerol, 4.1% SDS, 0.02% Bromophenol Blue, 5% β-mercaptoethanol) and boiled at 100°C for 10 mins. To blot for 3xFLAG-PilU or RpoA, 2 µL of each sample was electrophoretically separated on 10% SDS-PAGE gels. Following SDS-PAGE, the proteins were transferred to a polyvinylidene difluoride (PVDF) membrane by electrophoresis. Next, the membranes were blocked with 5% milk solution for 1 hr rocking at room temperature. To blot for 3xFLAG-pilU, primary mouse monoclonal anti-FLAG M2 antibody was added to the membrane. To blot for RpoA, primary mouse monoclonal anti-RpoA antibody was added, and membranes were incubated rocking at room temperature overnight. The next day, membranes were washed and incubated with anti-mouse horseradish peroxidase-conjugated secondary antibody for two hours rocking at room temperature. Lastly, blots were washed, developed with Pierce ECL Western blotting substrate, and imaged with a ProteinSimple FluorChem R system.

### Summary Statistics

See **Dataset S1** for summary statistics and a comprehensive list of statistical comparisons for all quantitative data in this manuscript.

## ACKNOLWEDGEMENTS

We would like to thank Marc Morais for helpful discussions. This work was supported by grant R35GM128674 from the National Institutes of Health (NIH) to ABD and R01GM148586 from NIH to JCG.

---

**Fig. S1.**
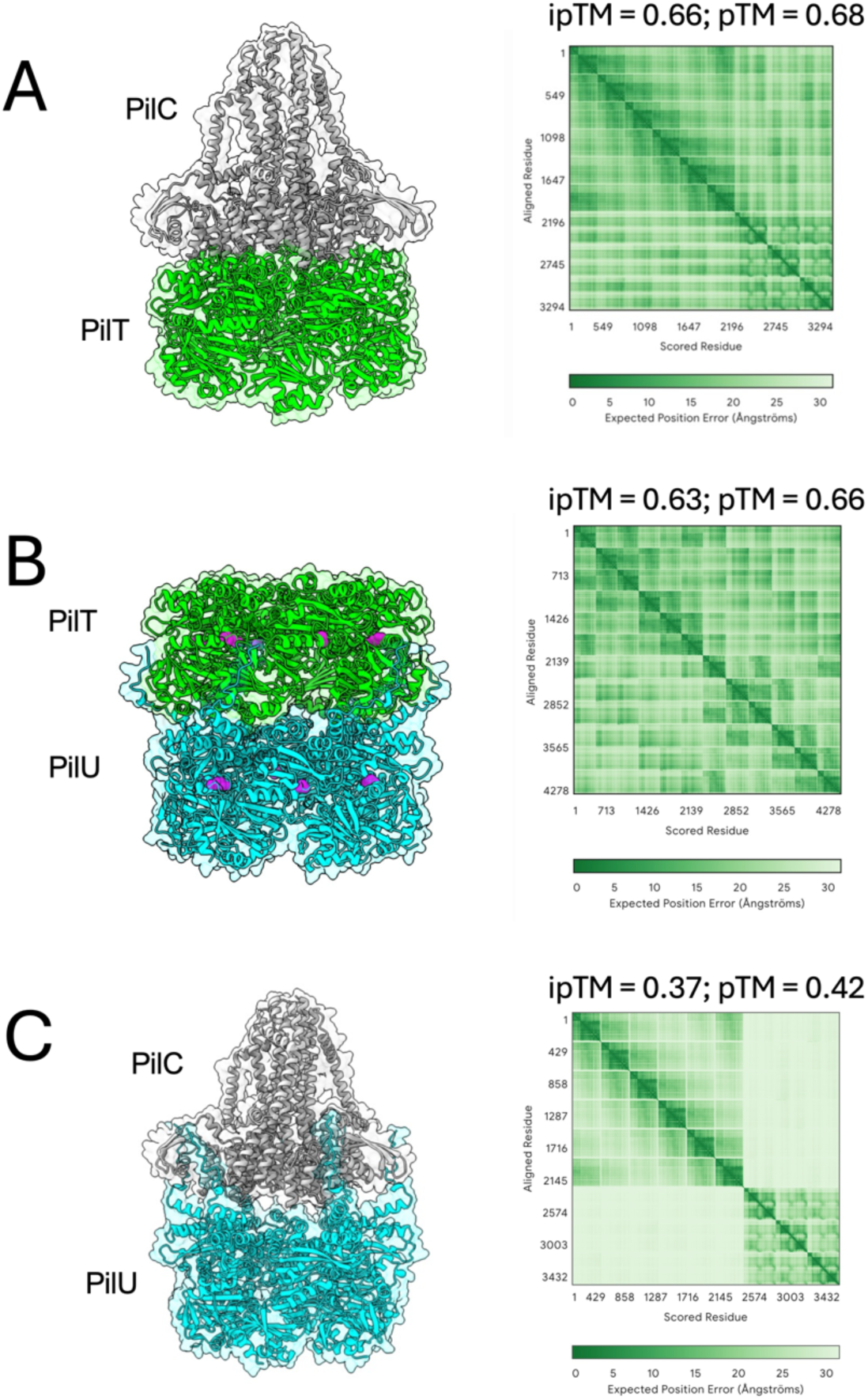
AF3 models of PilT-PilC, PilT-PilU, and PilU-PilC interactions. AF3 models (left) of the indicated *V. cholerae* proteins with their respective PAE maps and confidence scores (right). (**A**) PilT(x6)-PilC(x3) and (**B**) PilT(x6)-PilU(x6) were used to generate the PilC-PilT-PilU hybrid complex shown in Fig. 1A. The magenta spheres in **B** denote the Walker A residues in PilT and PilU (PilT^K136A^ and PilU^K134A^, respectively). (**C**) The PilU(x6)-PilC(x3) AF3 model had very low confidence metrics, suggesting that these proteins may not directly interact.

**Fig. S2.**
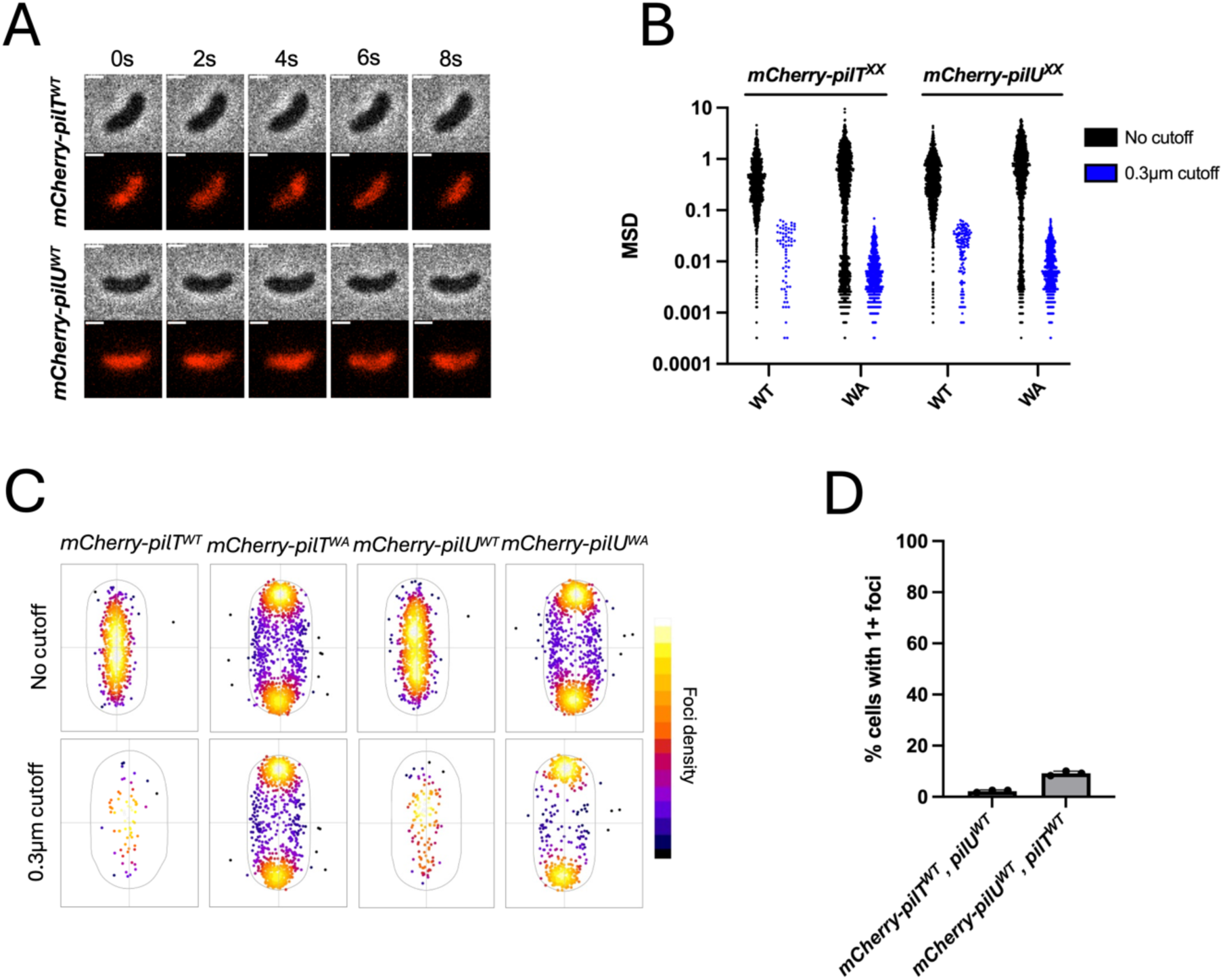
*mCherry-pilT^WA^* and *mCherry-pilU^WA^* form foci at the cell periphery. (**A**) Representative montage of timelapse imaging for the indicated strains highlights the lack of fluorescent foci in these backgrounds. Scale bars, 1 µM. (**B**) Mean-squared displacement (MSD) values of motor foci in the indicated strains. Black scatter plots represent MSD data collected with no cutoff for the distance foci can move between frames of the 10 s timelapse. Blue scatter plots represent MSD data collected for the same strains with a 0.3 µm cutoff for foci movement between frames. (**C**) Scatter plots of the localization of fluorescent foci in the same strains from **B** with and without a 0.3 µm distance cutoff. (**D**) Motor localization assays were performed on the indicated *V. cholerae* strains. For each strain, the percentage of cells (*n* = 120 cells analyzed) with at least one fluorescent focus that moves less than 0.3 µm between frames was determined. Data are from three independent biological replicates and are shown as the mean ± SD.

**Fig. S3.**
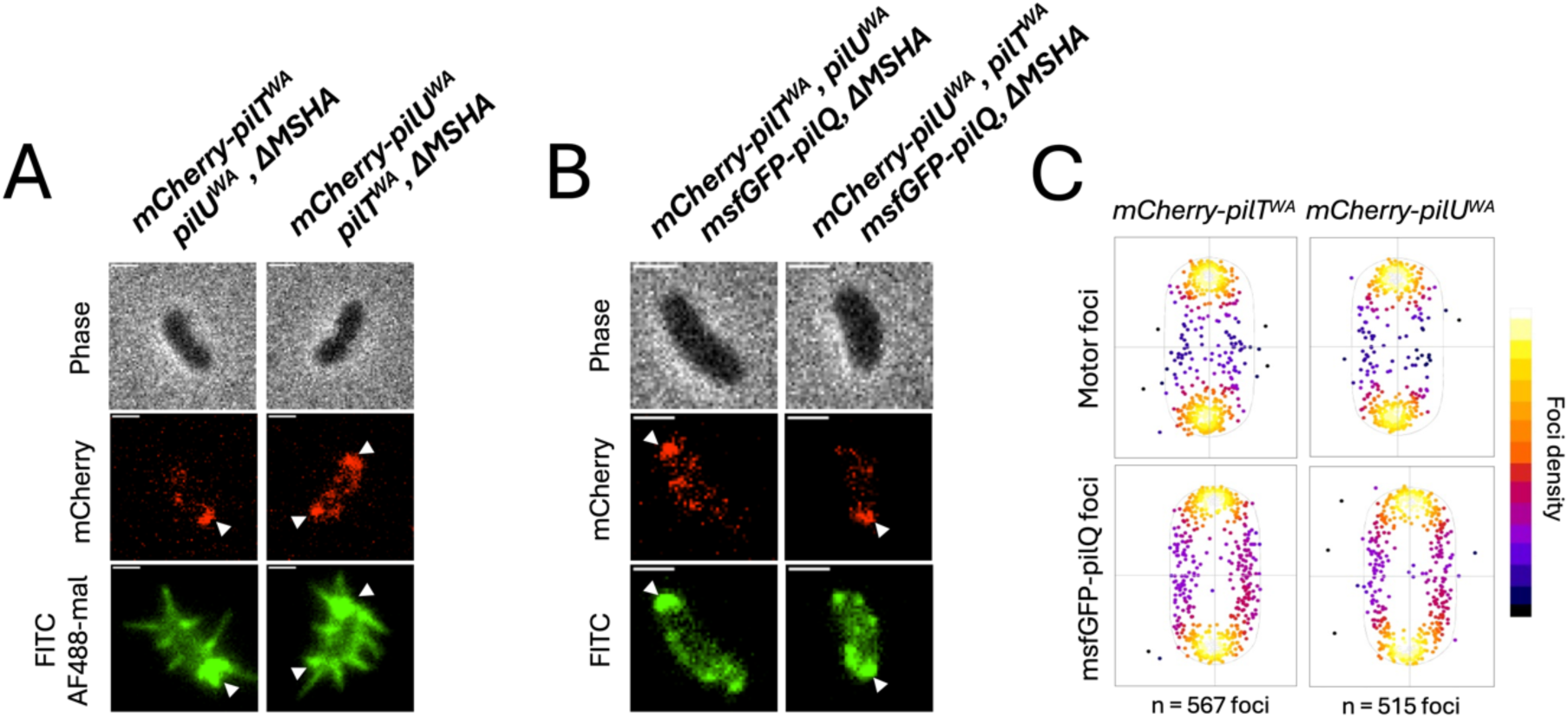
PilT and PilU foci colocalize with competence T4P machines. (**A**) *ΔMSHA* cells expressing *mCherry- pilT^WA^*or *mCherry-pilU^WA^*as indicated were labeled with AF488-mal to colocalize motor foci and competence pili. White carats denote motor focus localization at the base of competence T4P filaments. Scale bar, 1 µm. (**B)** *ΔMSHA* cells expressing *msfGFP-pilQ* and *mCherry-pilT^WA^*or *mCherry-pilU^WA^*as indicated. The representative images show motor foci colocalizing with msfGFP-PilQ foci (mCherry-PilT^WA^ and mCherry-PilU^WA^ foci colocalize with msfGFP-PilQ foci at a frequency of 99.3 ± 0.7% and 98.1% ± 1%, respectively). Scale bar, 1 µm. (**C**) Scatter heat maps denote the localization of motor foci and PilQ foci from the experiments described in **B**. These heat maps highlight that both PilT/PilU motor foci and PilQ foci are biased towards the cell poles. Data in **A-C** are representative of *n* = 3 biological replicates, and colocalization frequency is reported as the mean ± SD.

**Fig. S4.**
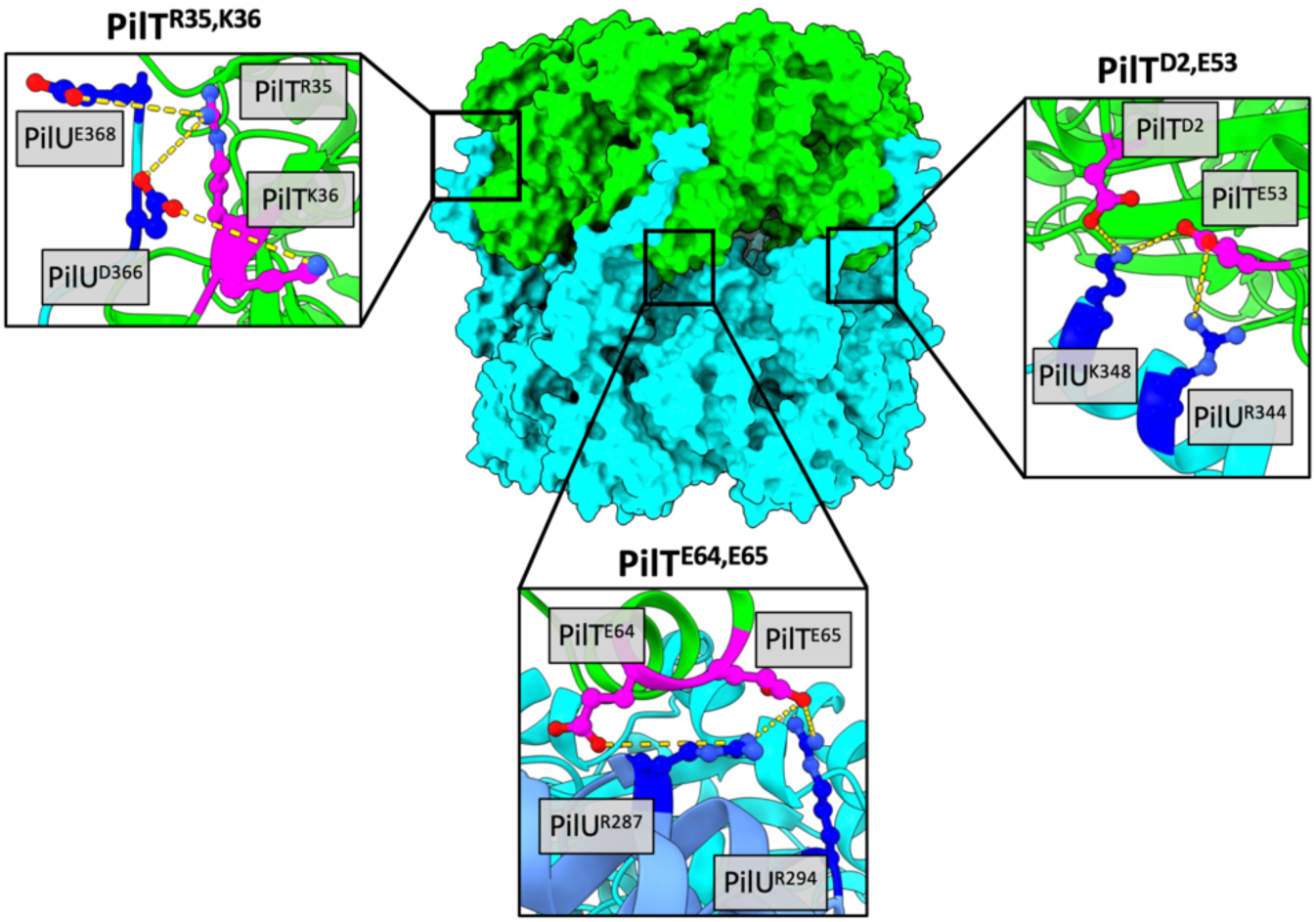
Contact analysis of the AF3 PilT-PilU model reveals three putative intermolecular salt bridges. AF3 model of the PilT (green)-PilU (cyan) complex. Insets show the residues involved in the three putative intermolecular salt bridges with PilT residues colored in magenta and PilU residues colored in blue.

**Fig. S5.**
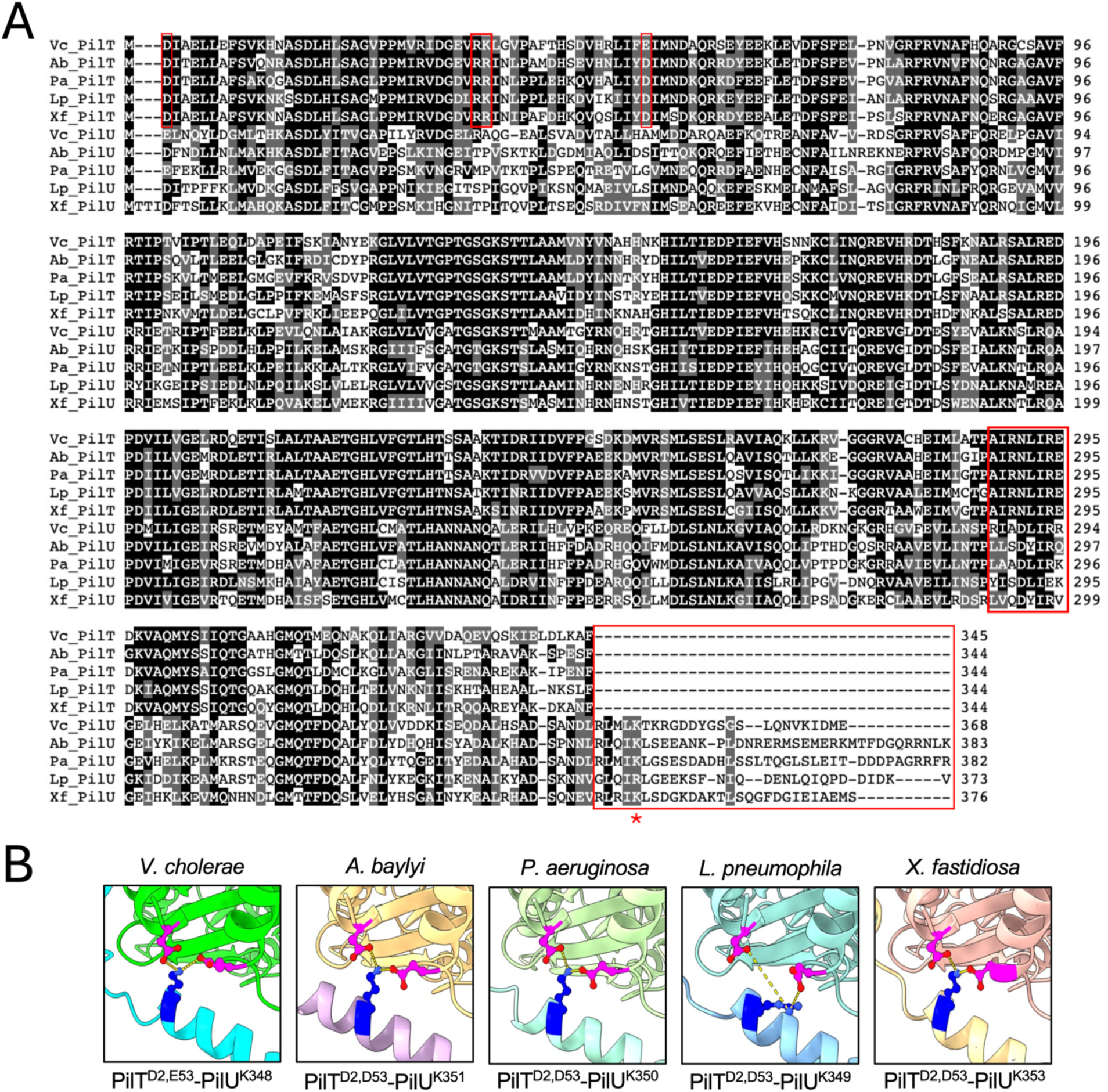
Sequence and structural alignments uncover conserved elements in PilT and PilU homologs. (**A**) Multiple sequence alignment of PilT and PilU homologs from the following bacterial species: Vc = *Vibrio cholerae*, Ab = *Acinetobacter baylyi*, Pa = *Pseudomonas aeruginosa*, Lp = *Legionella pneumophila*, and Xf = *Xylella fastidiosa*. Red boxes denote conservation of the PilT_Vc_ residues studied (D2, R35, K36, and E53), the PilT AIRNLIRE motif, and the extended C-terminus that is unique to PilU homologs. The red asterisk highlights conservation of the PilU_Vc_^K348^ residue among PilU homologs. (**B**) AF3 models of PilT-PilU complexes from the indicated species highlights the predicted conservation of the experimentally validated PilT_Vc_^D2,E53^-PilU_Vc_^K348^ intermolecular salt bridge.

**Fig. S6.**
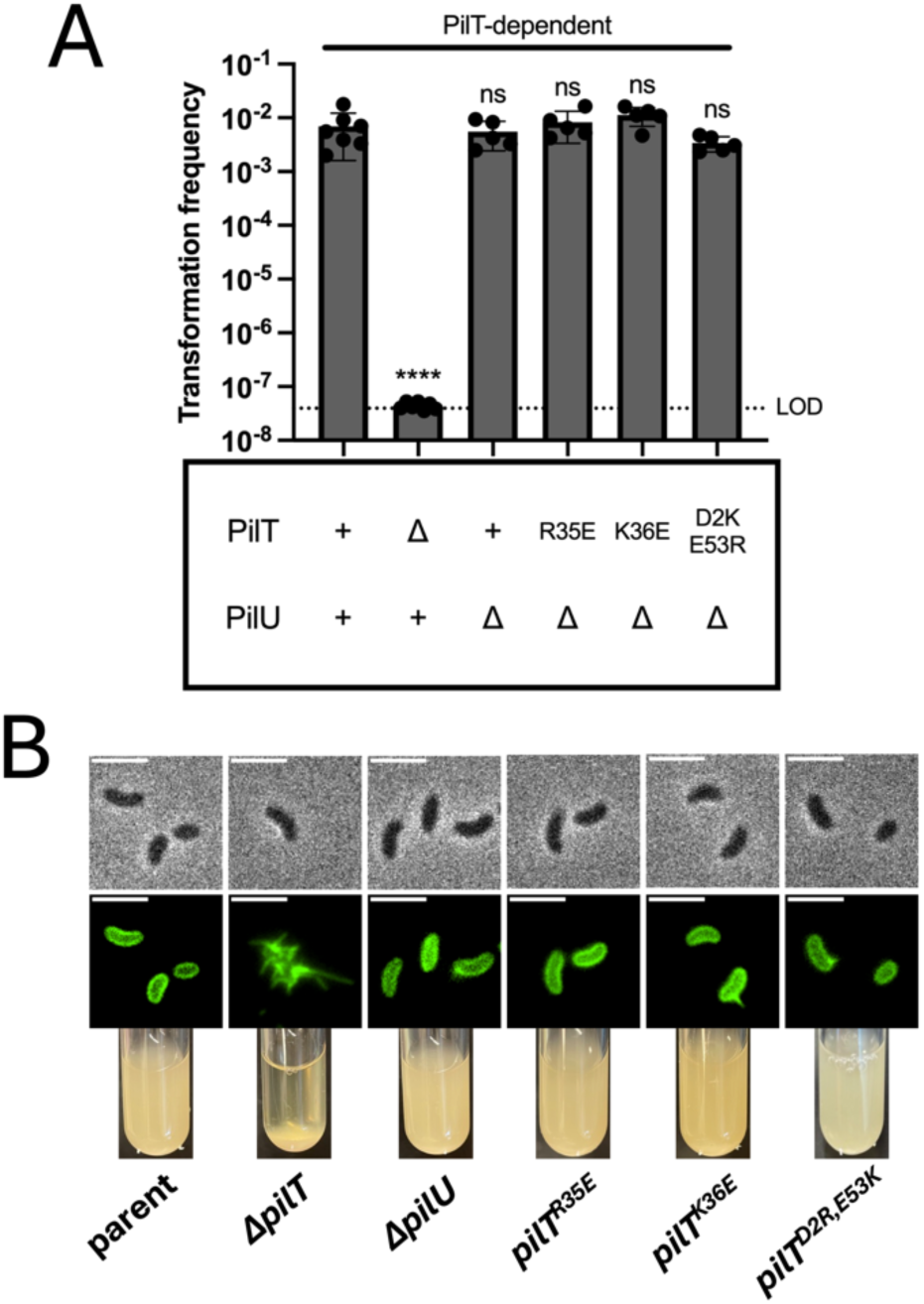
Mutating the PilT residues in the putative intermolecular salt bridges does not impact PilT-dependent retraction. (**A**) NT assays of the indicated *V. cholerae* strains demonstrates that mutating the residues involved in putative PilT-PilU intermolecular salt bridges does not impact PilT-dependent retraction (*i.e.*, in a background where PilT ATPase activity is intact). Data are from five independent biological replicates and shown as the mean ± SD. (**B**) Representative images of surface piliation (top panel) and cellular aggregation (bottom panel) for the indicated strains. Scale bar for micrographs, 3 µm. Data are representative of two independent experiments. Statistical comparisons were made by one-way ANOVA with Tukey’s multiple comparison test of the log-transformed data. ns, not significant; **** = *p* < 0.0001. Symbols directly above bars denote comparisons to the parent.

**Fig. S7.**
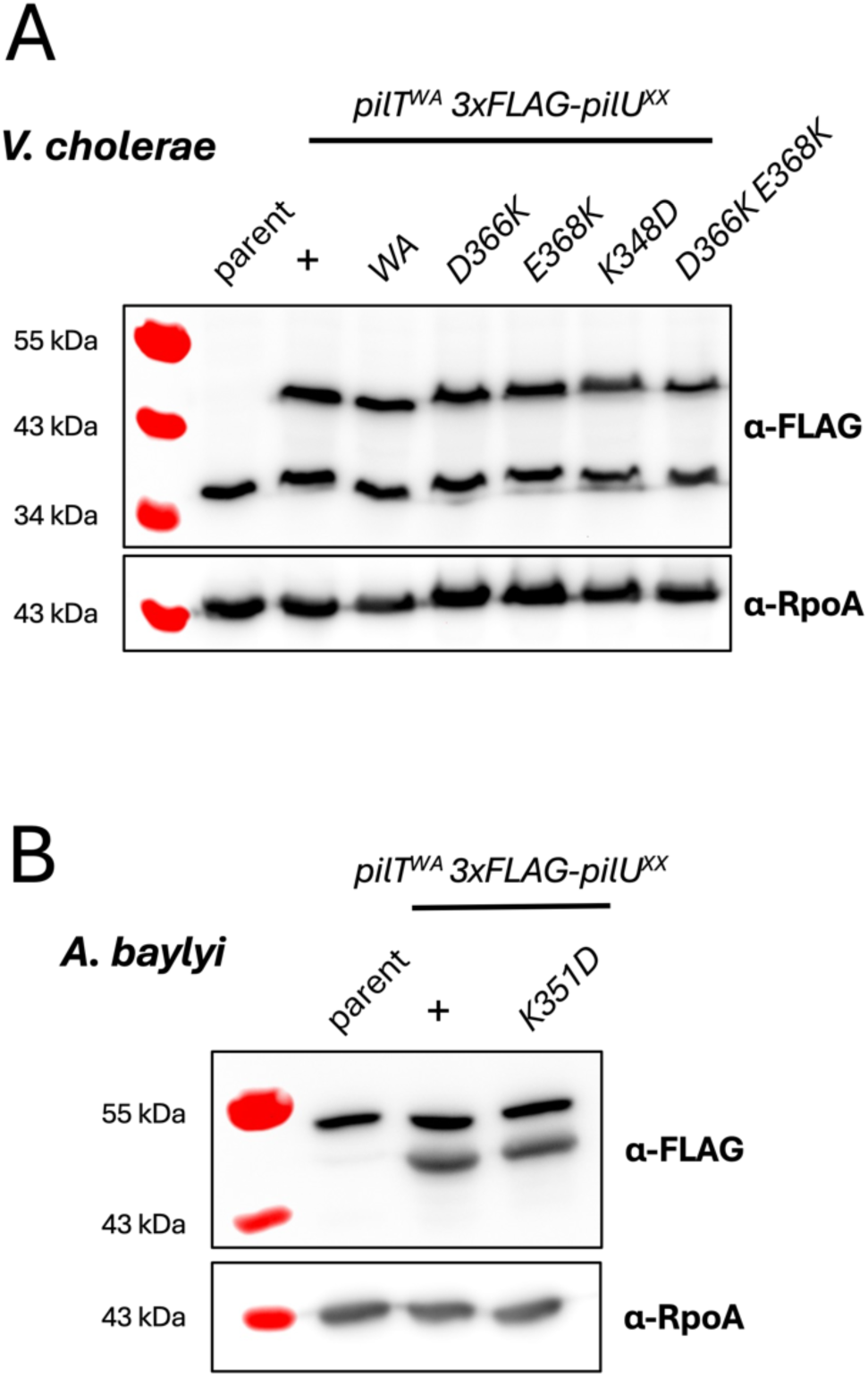
PilU point mutants do not impact PilU steady-state levels. Western blot analysis to assess steady-state levels of 3xFLAG-PilU and RpoA (loading control) from whole-cell lysates of the indicated point mutants in (**A**) *V. cholerae* and (**B**) *A. baylyi*. 3xFLAG-PilU is ∼45kDa in *V. cholerae* and ∼47kDa in *A. baylyi*. Notably, in anti-FLAG blots there is a distinct nonspecific band in both species (demarcated by the parent strain that lacks any FLAG-tagged constructs), which serves as an additional loading control. Blots in **A** and **B** are representative of three independent biological replicates.

**Fig. S8.**
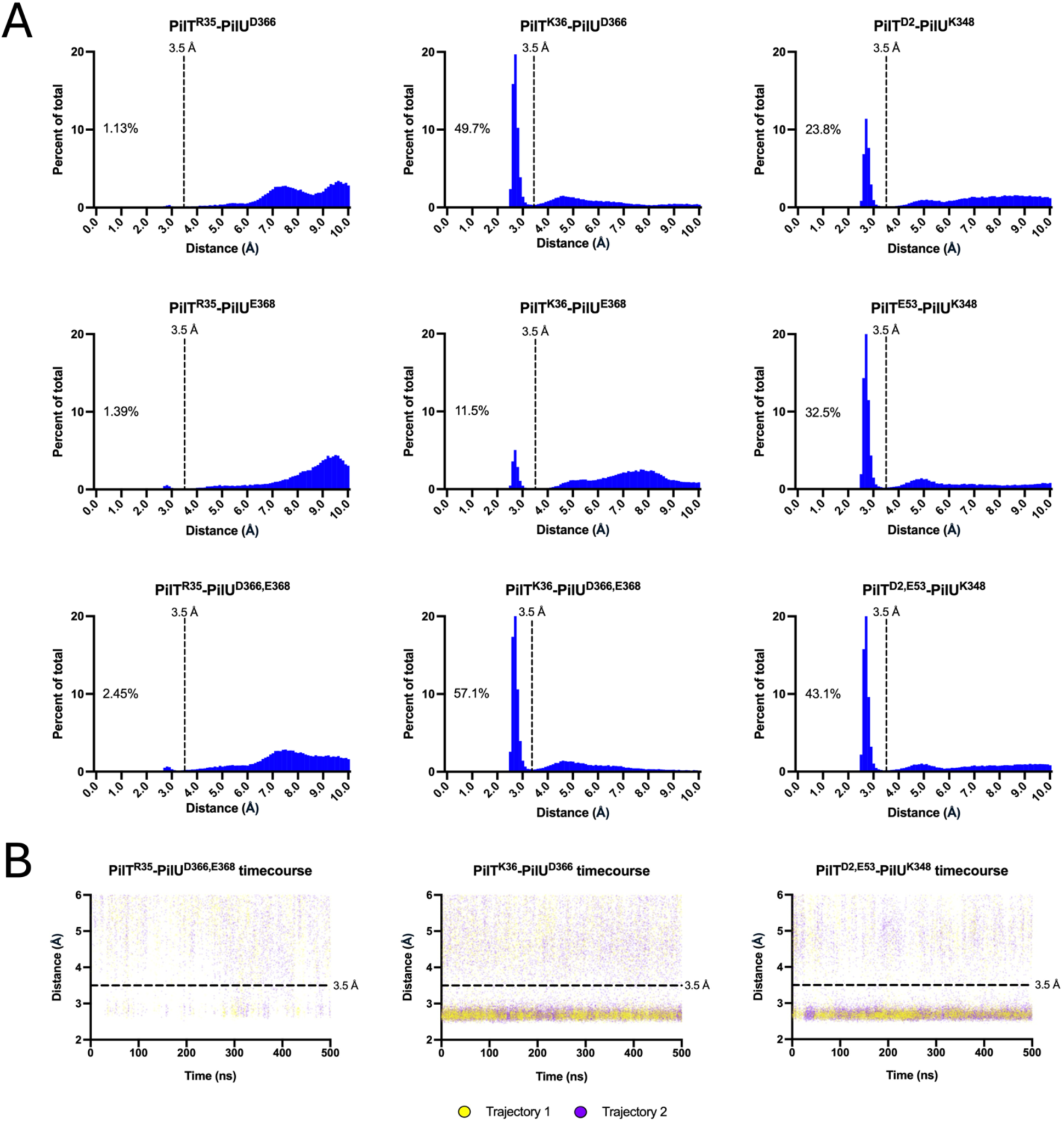
MD simulations of the AF3 PilT-PilU complex predict stable salt bridges between PilT^K36^-PilU^D366^ and PilT^D2,E53^-PilU^K348^. (**A**) Distance frequency histogram for the indicated residue pairs during the MD simulation of the AF3 PilT-PilU model. Distances between the side chains of the indicated residues were measured at 0.1-ns intervals over the 500-ns simulation. Percentage on graph denotes the amount of time that the indicated side chains are <3.5 Å (dotted line). Data are compiled from two independent MD simulations. (**B**) Time-course scatter plots of distance measurements for the indicated residue pairs. Trajectories 1 and 2 represent two independent replicates of the MD simulations.

**Fig. S9.**
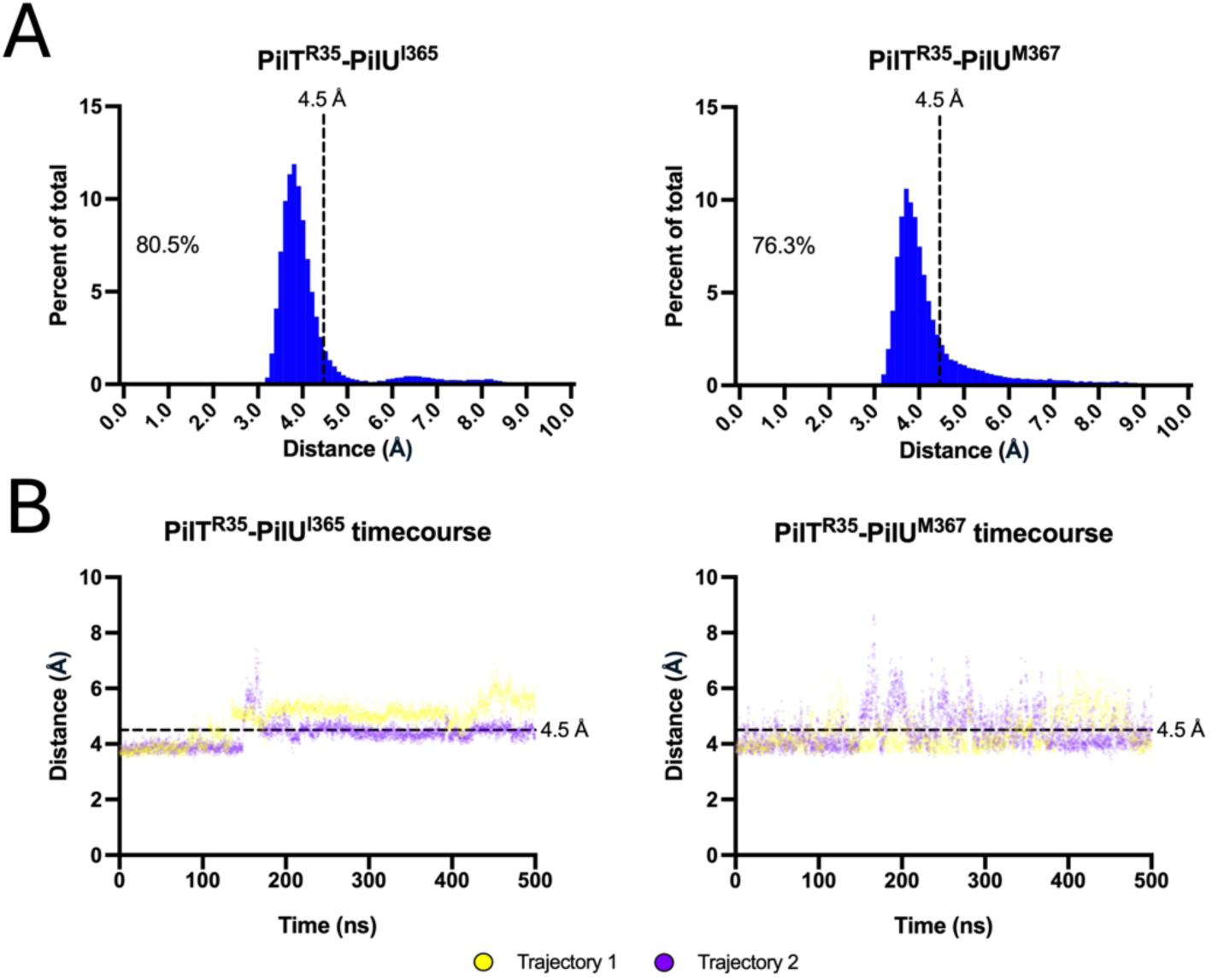
AF3-MD simulations support PilT^R35^-PilU^I365,M367^ interactions. (**A**) Distance frequency histograms of the indicated residues. Distances between the side chains of the indicated residues were recorded at 0.1-ns intervals over the 500-ns MD simulation. Percentages denote the amount of time that the residues were within 4.5 Å (dotted line). Data are compiled from two independent MD simulations. (**B**) Time-course scatter plots of the distance measurements for the indicated interactions. Trajectories 1 and 2 represent two independent replicates of the MD simulation.

**Fig. S10.**
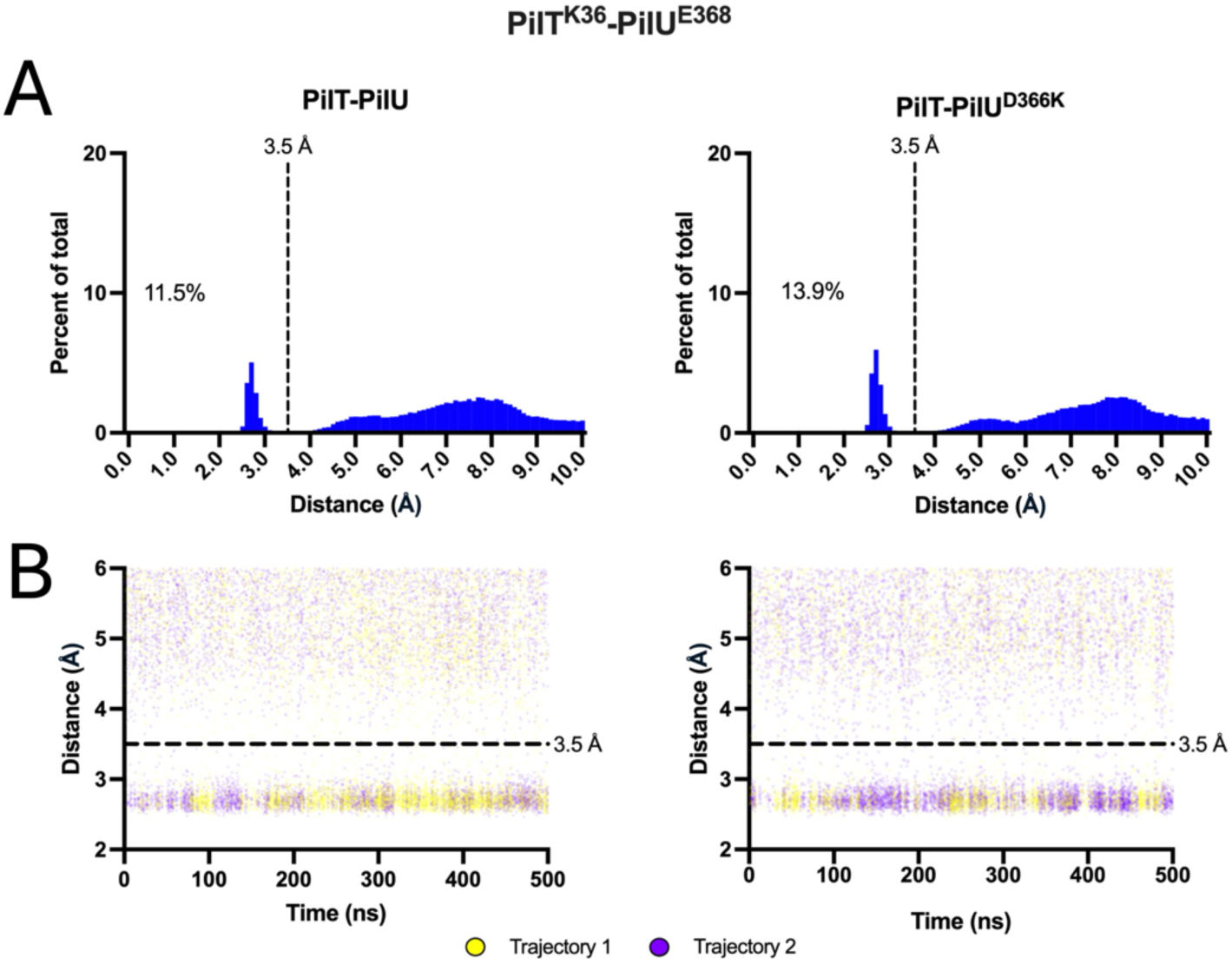
MD simulations predict intermediate contact between PilT^K36^-PilU^E368^ in the PilT-PilU and PilT- PilU^D366K^ AF3 complexes. (**A**) Distance frequency histograms of PilT^K36^-PilU^E368^ in the PilT-PilU vs PilT-PilU^D366K^ MD simulations as indicated. Distances between the side chains were measured at 0.1-ns intervals over the 500-ns simulation. Percentages on the histograms denote the amount of time the PilT^K36E^ and PilU^E368K^ are <3.5 Å apart (dotted line). Data are compiled from two independent MD simulations. (**B**) Timecourse scatter plots of the distances between PilT^K36E^-PilU^E368K^ in the PilT-PilU vs PilT-PilU^E368K^ MD simulations as indicated. Trajectories 1 and 2 represent two independent replicates of the MD simulation. In **A**, the data for the PilT-PilU complex is identical to the data presented in **Fig. S8** and are included here for ease of comparison.

**Fig. S11.**
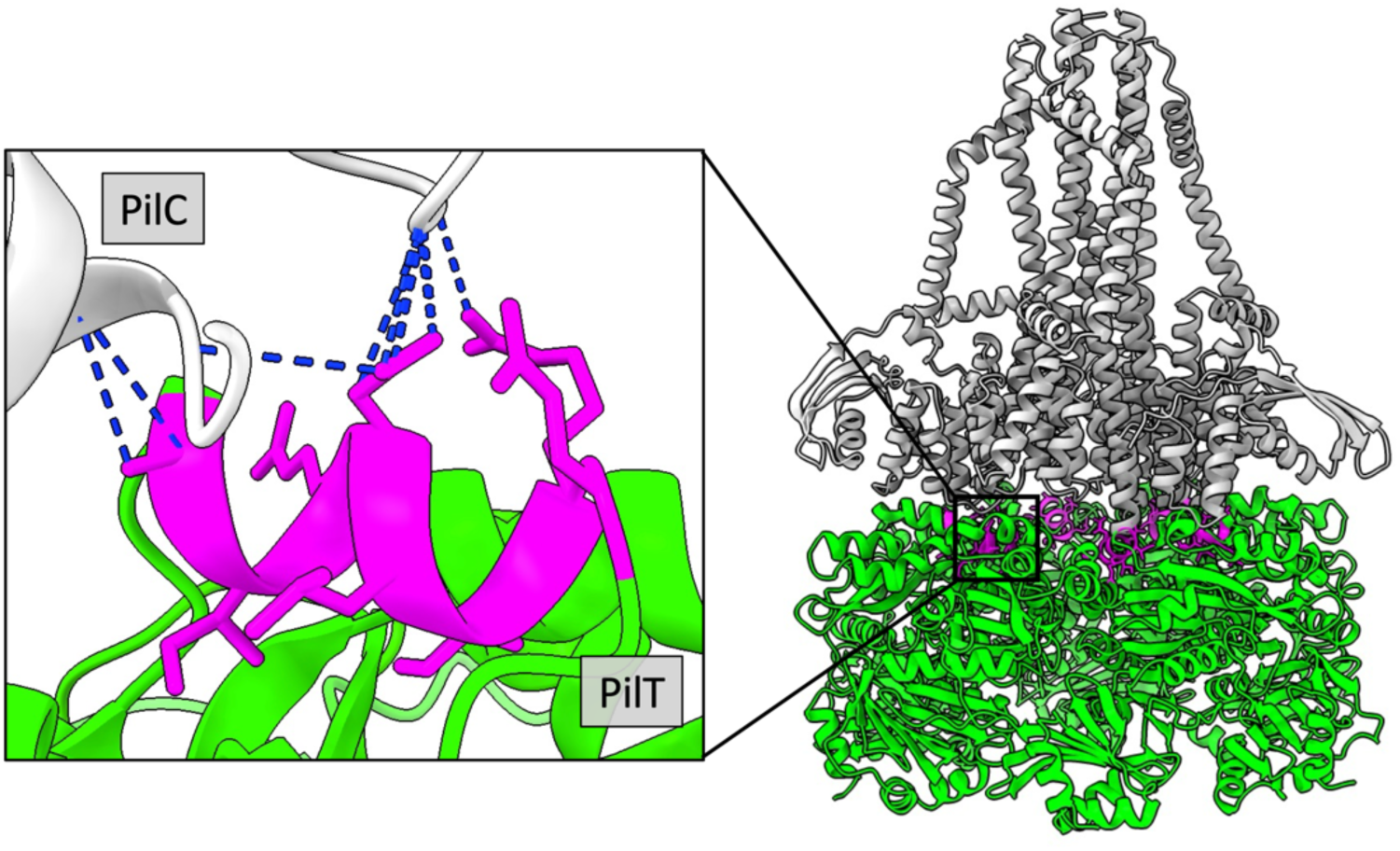
AF3 predicts that the AIRNLIRE motif is at the PilC-PilT interface. AF3 model of the PilC(3x)-PilT(6x) complex. Residues of the AIRNLIRE motif (PilT^A288-E295^) are colored in magenta. The inset shows a slice of the PilT- PilC model for a better view of the AIRNLIRE motif at the PilC-PilT interface. The blue dashed lines represent contacts predicted between the PilT AIRNLIRE motif and PilC.

**Table S1.** Strains used in this study.

| Strain # | Genotype | Figure | Use in manuscript | Name in manuscript |
| --- | --- | --- | --- | --- |
| AET249 / SAD4023 | E7946 Sm<sup>R</sup>, P<sub>tac</sub>-tfoX, ΔluxO, pilA<sup>S67C</sup>, ΔpilT::Smx<sup>R</sup> pilU<sup>K134A</sup>, ΔlacZ::Spec<sup>R</sup>-P<sub>pilT</sub>-mCherry-L5-pilT<sup>K136A</sup> | Fig 1b;<br>Fig 1c;<br>Fig S2 | Fluorescence microscopy foci quantification | mCherry-pilT<sup>WA</sup>, pilU<sup>WA</sup> |
| AET396 / SAD4024 | E7946 Sm<sup>R</sup>, P<sub>tac</sub>-tfoX, ΔluxO, pilA<sup>S67C</sup>, ΔpilT::Smx<sup>R</sup> pilU<sup>K134A</sup>, ΔlacZ::Spec<sup>R</sup>-P<sub>pilT</sub>-mCherry-L5-pilT<sup>K136A</sup>, ΔMSHA::Carb<sup>R</sup>, ΔpilC::Cm<sup>R</sup> | Fig 1c | Fluorescence microscopy foci quantification | mCherry-pilT<sup>WA</sup>, pilU<sup>WA</sup>, Δplatform |
| AET146 / SAD4025 | E7946 Sm<sup>R</sup>, P<sub>tac</sub>-tfoX, ΔluxO, pilA<sup>S67C</sup>, ΔlacZ::Spec<sup>R</sup>-P<sub>pilT</sub>-mCherry-L5-pilT<sup>K136A</sup>, ΔpilTU::Tm<sup>R</sup> | Fig 1c;<br>Fig 2e | Fluorescence microscopy foci quantification | mCherry-pilT<sup>WA</sup>, ΔpilU |
| AET222 / SAD4026 | E7946 Sm<sup>R</sup>, P<sub>tac</sub>-tfoX, ΔluxO, pilA<sup>S67C</sup>, ΔlacZ::Spec<sup>R</sup>-P<sub>pilT</sub>-mCherry-L5-pilT<sup>K136A</sup>, ΔpilTU::Tm<sup>R</sup>, ΔMSHA::Carb<sup>R</sup>, ΔpilC::Cm<sup>R</sup> | Fig 1c | Fluorescence microscopy foci quantification | mCherry-pilT<sup>WA</sup>, ΔpilU, Δplatform |
| AET159 / SAD4027 | E7946 Sm<sup>R</sup>, P<sub>tac</sub>-tfoX, ΔluxO, pilA<sup>S67C</sup>, ΔpilT::Smx<sup>R</sup> mCherry-L5-pilU<sup>K134A</sup>, ΔlacZ::Spec<sup>R</sup>-P<sub>pilT</sub>-pilT<sup>K136A</sup> | Fig 1b;<br>Fig 1d;<br>Fig 2d;<br>Fig S2 | Fluorescence microscopy foci quantification | mCherry-pilU<sup>WA</sup>, pilT<sup>WA</sup> |
| AET205 / SAD4028 | E7946 Sm<sup>R</sup>, P<sub>tac</sub>-tfoX, ΔluxO, pilA<sup>S67C</sup>, ΔpilT::Smx<sup>R</sup> mCherry-L5-pilU<sup>K134A</sup>, ΔlacZ::Spec<sup>R</sup>-P<sub>pilT</sub>-pilT<sup>K136A</sup>, ΔMSHA::Carb<sup>R</sup>, ΔpilC::Cm<sup>R</sup> | Fig 1d | Fluorescence microscopy foci quantification | mCherry-pilU<sup>WA</sup>, pilT<sup>WA</sup>, Δplatform |
| AET153 / SAD4029 | E7946 Sm<sup>R</sup>, P<sub>tac</sub>-tfoX, ΔluxO, pilA<sup>S67C</sup>, ΔpilT::Smx<sup>R</sup> mCherry-L5-pilU<sup>K134A</sup> | Fig 1d;<br>Fig 2d | Fluorescence microscopy foci quantification | mCherry-pilU<sup>WA</sup>, ΔpilT |
| AET155 / SAD4030 | E7946 Sm<sup>R</sup>, P<sub>tac</sub>-tfoX, ΔluxO, pilA<sup>S67C</sup>, ΔpilT::Smx<sup>R</sup> gfp-L5-pilU<sup>K134A</sup>, ΔlacZ::Spec<sup>R</sup>-P<sub>pilT</sub>-pilT<sup>K136A</sup>-L5-mCherry | Fig 1e;<br>Fig 1f | Fluorescence co-localization | pilT<sup>WA</sup>-mCherry, gfp-pilU<sup>WA</sup> |
| TND0905 / SAD2468 / (18, 26) | E7946 Sm<sup>R</sup>, P<sub>tac</sub>-tfoX, ΔluxO, pilA<sup>S67C</sup> | Fig 2b-c;<br>Fig S6;<br>Fig S7a | Transformation, pilus labeling, cellular aggregation assays, and western blots | Parent |
| AET111 / SAD4031 | E7946 Sm<sup>R</sup>, P<sub>tac</sub>-tfoX, ΔluxO, pilA<sup>S67C</sup>, ΔpilT::Smx<sup>R</sup> | Fig 2b-c;<br>Fig S6 | Transformation, pilus labeling, and cellular aggregation assays | ΔpilT |
| TND1515 / SAD4032 | E7946 Sm<sup>R</sup>, P<sub>tac</sub>-tfoX, ΔluxO, pilA<sup>S67C</sup>, ΔpilU::Tm<sup>R</sup> | Fig 2b-c;<br>Fig S6 | Transformation, pilus labeling, and cellular aggregation assays | ΔpilU |
| TND4516 / SAD4033 | E7946 Sm<sup>R</sup>, P<sub>tac</sub>-tfoX, ΔluxO, pilA<sup>S67C</sup>, ΔpilT::Smx<sup>R</sup>, ΔlacZ::Spec<sup>R</sup>-P<sub>pilT</sub>-pilT<sup>K136A</sup> | Fig 2b-c;<br>Fig 3b-c | Transformation, pilus labeling, and cellular aggregation assays | pilT<sup>WA</sup> |
| TND4360 / SAD4034 | E7946 Sm<sup>R</sup>, P<sub>tac</sub>-tfoX, ΔluxO, pilA<sup>S67C</sup>, ΔpilTU::Tm<sup>R</sup>, ΔlacZ::Spec<sup>R</sup>-P<sub>pilT</sub>-pilT<sup>K136A</sup> | Fig 2b-c;<br>Fig 3b-c | Transformation, pilus labeling, and cellular aggregation assays | pilT<sup>WA</sup>, ΔpilU |
| TND4361 / SAD4035 | E7946 Sm<sup>R</sup>, P<sub>tac</sub>-tfoX, ΔluxO, pilA<sup>S67C</sup>, ΔpilTU::Tm<sup>R</sup>, ΔlacZ::Spec<sup>R</sup>-P<sub>pilT</sub>-pilT<sup>E64A,E65A</sup> | Fig 2b-c | Transformation, pilus labeling, and cellular aggregation assays | pilT<sup>E64A,E65A</sup> |
| TND4362 / SAD4036 | E7946 Sm<sup>R</sup>, P<sub>tac</sub>-tfoX, ΔluxO, pilA<sup>S67C</sup>, ΔpilTU::Tm<sup>R</sup>, ΔlacZ::Spec<sup>R</sup>-P<sub>pilT</sub>-pilT<sup>R35A,K36A</sup> | Fig 2b-c | Transformation, pilus labeling, and cellular aggregation assays | pilT<sup>R35A,K36A</sup> |
| AET265 / SAD4037 | E7946 Sm<sup>R</sup>, P<sub>tac</sub>-tfoX, ΔluxO, pilA<sup>S67C</sup>, ΔpilTU::Tm<sup>R</sup>, ΔlacZ::Spec<sup>R</sup>-P<sub>pilT</sub>-pilT<sup>D2A,E53A</sup> | Fig 2b-c | Transformation, pilus labeling, and cellular aggregation assays | pilT<sup>D2A,E53A</sup> |
| AET112 / SAD4038 | E7946 Sm<sup>R</sup>, P<sub>tac</sub>-tfoX, ΔluxO, pilA<sup>S67C</sup>, ΔpilT::Smx<sup>R</sup>, ΔlacZ::Spec<sup>R</sup>-P<sub>pilT</sub>-pilT<sup>E64A,E65A,K136A</sup> | Fig 2b-c | Transformation, pilus labeling, and cellular aggregation assays | pilT<sup>WA,E64A,E65A</sup> |
| AET113 / SAD4039 | E7946 Sm<sup>R</sup>, P<sub>tac</sub>-tfoX, ΔluxO, pilA<sup>S67C</sup>, ΔpilT::Smx<sup>R</sup>, ΔlacZ::Spec<sup>R</sup>-P<sub>pilT</sub>-pilT<sup>R35A,K36A,K136A</sup> | Fig 2b-c | Transformation, pilus labeling, and cellular aggregation assays | pilT<sup>WA,R35A,K36A</sup> |
| AET275 / SAD4040 | E7946 Sm<sup>R</sup>, P<sub>tac</sub>-tfoX, ΔluxO, pilA<sup>S67C</sup>, ΔpilT::Smx<sup>R</sup>, ΔlacZ::Spec<sup>R</sup>-P<sub>pilT</sub>-pilT<sup>D2A,E53A,K136A</sup> | Fig 2b-c | Transformation, pilus labeling, and cellular aggregation assays | pilT<sup>WA,D2A,E53A</sup> |
| AET262 / SAD4041 | E7946 Sm<sup>R</sup>, P<sub>tac</sub>-tfoX, ΔluxO, pilA<sup>S67C</sup>, ΔpilT::Smx<sup>R</sup> mCherry-L5-pilU<sup>K134A</sup>, ΔlacZ::Spec<sup>R</sup>-P<sub>pilT</sub>-pilT<sup>E64A,E65A,K136A</sup> | Fig 2d | Fluorescence microscopy foci quantification | mCherry-pilU<sup>WA</sup>, pilT<sup>WA,E64A,E65A</sup> |
| AET263 / SAD4042 | E7946 Sm<sup>R</sup>, P<sub>tac</sub>-tfoX, ΔluxO, pilA<sup>S67C</sup>, ΔpilT::Smx<sup>R</sup> mCherry-L5-pilU<sup>K134A</sup>, ΔlacZ::Spec<sup>R</sup>-P<sub>pilT</sub>-pilT<sup>R35A,K36A,K136A</sup> | Fig 2d | Fluorescence microscopy foci quantification | mCherry-pilU<sup>WA</sup>, pilT<sup>WA,R35A,K36A</sup> |
| AET277 / SAD4043 | E7946 Sm<sup>R</sup>, P<sub>tac</sub>-tfoX, ΔluxO, pilA<sup>S67C</sup>, ΔpilT::Smx<sup>R</sup> mCherry-L5-pilU<sup>K134A</sup>, ΔlacZ::Spec<sup>R</sup>-P<sub>pilT</sub>-pilT<sup>D2A,E53A,K136A</sup> | Fig 2d | Fluorescence microscopy foci quantification | mCherry-pilU<sup>WA</sup>, pilT<sup>WA,D2A,E53A</sup> |
| AET271 / SAD4044 | E7946 Sm<sup>R</sup>, P<sub>tac</sub>-tfoX, ΔluxO, pilA<sup>S67C</sup>, ΔpilTU::Tm<sup>R</sup>, ΔlacZ::Spec<sup>R</sup>-P<sub>pilT</sub>-mCherry-L5-pilT<sup>E64A,E65A,K136A</sup> | Fig 2e | Fluorescence microscopy foci quantification | mCherry-pilT<sup>WA,E64A,E65A</sup> |
| AET270 / SAD4045 | E7946 Sm<sup>R</sup>, P<sub>tac</sub>-tfoX, ΔluxO, pilA<sup>S67C</sup>, ΔpilTU::Tm<sup>R</sup>, ΔlacZ::Spec<sup>R</sup>-P<sub>pilT</sub>-mCherry-L5-pilT<sup>R35A,K36A,K136A</sup> | Fig 2e | Fluorescence microscopy foci quantification | mCherry-pilT<sup>WA,R35A K36A</sup> |
| AET278 / SAD4046 | E7946 Sm<sup>R</sup>, P<sub>tac</sub>-tfoX, ΔluxO, pilA<sup>S67C</sup>, ΔpilTU::Tm<sup>R</sup>, ΔlacZ::Spec<sup>R</sup>-P<sub>pilT</sub>-mCherry-L5-pilT<sup>D2A,E53A,K136A</sup> | Fig 2e | Fluorescence microscopy foci quantification | mCherry-pilT<sup>WA,D2A,E53A</sup> |
| AET189 / SAD4047 | E7946 Sm<sup>R</sup>, P<sub>tac</sub>-tfoX, ΔluxO, pilA<sup>S67C</sup>, ΔpilT::Smx<sup>R</sup>, ΔlacZ::Spec<sup>R</sup>-P<sub>pilT</sub>-pilT<sup>R35E,K136A</sup> | Fig 3b-c | Transformation, pilus labeling, and cellular aggregation assays | pilT<sup>WA,R35E</sup> |
| TND4578 / SAD4048 | E7946 Sm<sup>R</sup>, P<sub>tac</sub>-tfoX, ΔluxO, pilA<sup>S67C</sup>, ΔpilT::Smx<sup>R</sup>, pilU<sup>D366K,E368K</sup>, ΔlacZ::Spec<sup>R</sup>-P<sub>pilT</sub>-pilT<sup>K136A</sup> | Fig 3b-c | Transformation, pilus labeling, and cellular aggregation assays | pilT<sup>WA</sup>, pilU<sup>D366K,E368K</sup> |
| AET391 / SAD4049 | E7946 Sm<sup>R</sup>, P<sub>tac</sub>-tfoX, ΔluxO, pilA<sup>S67C</sup>, ΔpilT::Smx<sup>R</sup>, pilU<sup>D366K,E368K</sup>, ΔlacZ::Spec<sup>R</sup>-P<sub>pilT</sub>-pilT<sup>R35E,K136A</sup> | Fig 3b-c | Transformation, pilus labeling, and cellular aggregation assays | pilT<sup>WA,R35E</sup>, pilU<sup>D366K,E368K</sup> |
| AET190 / SAD4050 | E7946 Sm<sup>R</sup>, P<sub>tac</sub>-tfoX, ΔluxO, pilA<sup>S67C</sup>, ΔpilT::Smx<sup>R</sup>, ΔlacZ::Spec<sup>R</sup>-P<sub>pilT</sub>-pilT<sup>K36E,K136A</sup> | Fig 3b-c | Transformation, pilus labeling, and cellular aggregation assays | pilT<sup>WA,K36E</sup> |
| AET200 / SAD4051 | E7946 Sm<sup>R</sup>, P<sub>tac</sub>-tfoX, ΔluxO, pilA<sup>S67C</sup>, ΔpilT::Smx<sup>R</sup>, pilU<sup>D366K</sup>, ΔlacZ::Spec<sup>R</sup>-P<sub>pilT</sub>-pilT<sup>K136A</sup> | Fig 3b-c | Transformation, pilus labeling, and cellular aggregation assays | pilT<sup>WA</sup>, pilU<sup>D366K</sup> |
| AET201 / SAD4052 | E7946 Sm<sup>R</sup>, P<sub>tac</sub>-tfoX, ΔluxO, pilA<sup>S67C</sup>, ΔpilT::Smx<sup>R</sup>, pilU<sup>E368K</sup>, ΔlacZ::Spec<sup>R</sup>-P<sub>pilT</sub>-pilT<sup>K136A</sup> | Fig 3b-c | Transformation, pilus labeling, and cellular aggregation assays | pilT<sup>WA</sup>, pilU<sup>E368K</sup> |
| AET198 / SAD4053 | E7946 Sm<sup>R</sup>, P<sub>tac</sub>-tfoX, ΔluxO, pilA<sup>S67C</sup>, ΔpilT::Smx<sup>R</sup>, pilU<sup>D366K</sup>, ΔlacZ::Spec<sup>R</sup>-P<sub>pilT</sub>-pilT<sup>K36E,K136A</sup> | Fig 3b-c | Transformation, pilus labeling, and cellular aggregation assays | pilT<sup>WA,K36E</sup>, pilU<sup>D366K</sup> |
| AET199 / SAD4054 | E7946 Sm<sup>R</sup>, P<sub>tac</sub>-tfoX, ΔluxO, pilA<sup>S67C</sup>, ΔpilT::Smx<sup>R</sup>, pilU<sup>E368K</sup>, ΔlacZ::Spec<sup>R</sup>-P<sub>pilT</sub>-pilT<sup>K36E,K136A</sup> | Fig 3b-c | Transformation, pilus labeling, and cellular aggregation assays | pilT<sup>WA,K36E</sup>, pilU<sup>E368K</sup> |
| AET475 / SAD4106 | E7946 Sm<sup>R</sup>, P<sub>tac</sub>-tfoX, ΔluxO, pilA<sup>S67C</sup>, ΔpilT::Smx<sup>R</sup>, pilU<sup>D366K,E368K</sup>, ΔlacZ::Spec<sup>R</sup>-P<sub>pilT</sub>-pilT<sup>K36E,K136A</sup> | Fig 3b-c | Transformation, pilus labeling, and cellular aggregation assays | pilT<sup>WA,K36E</sup>, pilU<sup>D366K,E368K</sup> |
| AET345 / SAD4055 | E7946 Sm<sup>R</sup>, P<sub>tac</sub>-tfoX, ΔluxO, pilA<sup>S67C</sup>, ΔpilT::Smx<sup>R</sup>, ΔlacZ::Spec<sup>R</sup>-P<sub>pilT</sub>-pilT<sup>D2K,E53R,K136A</sup> | Fig 3b-c | Transformation, pilus labeling, and cellular aggregation assays | pilT<sup>WA,D2K,E53R</sup> |
| AET337 / SAD4056 | E7946 Sm<sup>R</sup>, P<sub>tac</sub>-*tfoX*, $\Delta luxO$, *pilA*<sup>S67C</sup>, $\Delta pilT::Smx^R$ *pilU*<sup>K348D</sup>, $\Delta lacZ::Spec^R$-P<sub>*pilT*</sub>-*pilT*<sup>K136A</sup> | Fig 3b-c | Transformation, pilus labeling, and cellular aggregation assays | *pilT*<sup>WA</sup>, *pilU*<sup>K348D</sup> |
| AET358 / SAD4057 | E7946 Sm<sup>R</sup>, P<sub>tac</sub>-*tfoX*, $\Delta luxO$, *pilA*<sup>S67C</sup>, $\Delta pilT::Smx^R$ *pilU*<sup>K348D</sup>, $\Delta lacZ::Spec^R$-P<sub>*pilT*</sub>-*pilT*<sup>D2K,E53R,K136A</sup> | Fig 3b-c | Transformation, pilus labeling, and cellular aggregation assays | *pilT*<sup>WA,D2K,E53R</sup>, *pilU*<sup>K348D</sup> |
| TND3433 / SAD4058 / (24) | $\Delta VC1807::Cm^R$-P<sub>tac</sub>-*gfp::parST1*, $\Delta lacZ::Spec^R$-P<sub>tac</sub>-*mKate2* | Fig 4a-c | Chitin biofilm natural transformation assays | Parent (chitin biofilm NT assays) |
| TND4366 / SAD4059 / (24) | $\Delta VC1807::Cm^R$-P<sub>tac</sub>-*gfp::parST1*, $\Delta lacZ::Spec^R$-P<sub>tac</sub>-*mKate2*, $\Delta pilU::Tm^R$ | Fig 4a-c | Chitin biofilm natural transformation assays | $\Delta pilU$ (chitin biofilm NT assays) |
| AET254 / SAD4060 | $\Delta VC1807::Cm^R$-P<sub>tac</sub>-*gfp::parST1*, $\Delta lacZ::Spec^R$-P<sub>tac</sub>-*mKate2*, *pilT*<sup>R35E</sup> | Fig 4a-c | Chitin biofilm natural transformation assays | *pilT*<sup>R35E</sup> (chitin biofilm NT assays) |
| AET401 / SAD4107 | $\Delta VC1807::Cm^R$-P<sub>tac</sub>-*gfp::parST1*, $\Delta lacZ::Spec^R$-P<sub>tac</sub>-*mKate2*, *pilT*<sup>K136A</sup> | Fig 4a-c | Chitin biofilm natural transformation assays | *pilT*<sup>K136A</sup> (chitin biofilm NT assays) |
| AET402 / SAD4108 | $\Delta VC1807::Cm^R$-P<sub>tac</sub>-*gfp::parST1*, $\Delta lacZ::Spec^R$-P<sub>tac</sub>-*mKate2*, *pilU*<sup>K134A</sup> | Fig 4a-c | Chitin biofilm natural transformation assays | *pilU*<sup>K134A</sup> (chitin biofilm NT assays) |
| TND0428 / SAD2537 / (18) | *comP*<sup>T129C</sup> | Fig 5b; Fig S7b | *A. baylyi* natural transformation assays and Western blots | Parent (*A. baylyi* NT assays and Westerns) |
| TND0799 / SAD2544 / (18) | *comP*<sup>T129C</sup>, $\Delta pilTU::Zeo^R$ | Fig 5b | *A. baylyi* natural transformation assays | $\Delta pilT$ (*A. baylyi* NT assays) |
| TND0798 / SAD2543 / (18) | *comP*<sup>T129C</sup>, $\Delta pilU::Zeo^R$ | Fig 5b | *A. baylyi* natural transformation assays | $\Delta pilU$ (*A. baylyi* NT assays) |
| AET392 / SAD4061 | *comP*<sup>T129C</sup>, *pilT*<sup>D2K,D53R</sup>, $\Delta pilU::Zeo^R$ | Fig 5b | *A. baylyi* natural transformation assays | *pilT*<sup>D2K,D53R</sup> (*A. baylyi* NT assays) |
| TND1122 / SAD2540 / (18) | *comP*<sup>T129C</sup>, *pilT*<sup>K136A</sup> | Fig 5b | *A. baylyi* natural transformation assays | *pilT*<sup>WA</sup> (*A. baylyi* NT assays) |
| TND1135 / SAD2538 / (18) | *comP*<sup>T129C</sup>, *pilT*<sup>K136A</sup> Δ*pilU*::Zeo<sup>R</sup> | Fig 5b | *A. baylyi* natural transformation assays | *pilT*<sup>WA</sup>, Δ*pilU* (*A. baylyi* NT assays) |
| AET316 / SAD4062 | *comP*<sup>T129C</sup>, *pilT*<sup>D2K,D53R,K136A</sup> | Fig 5b | *A. baylyi* natural transformation assays | *pilT*<sup>WA,D2K,D53R</sup> (*A. baylyi* NT assays) |
| AET302 / SAD4063 | *comP*<sup>T129C</sup>, *pilT*<sup>K136A</sup> *pilU*<sup>K351D</sup> | Fig 5b | *A. baylyi* natural transformation assays | *pilT*<sup>WA</sup>, *pilU*<sup>K351D</sup> (*A. baylyi* NT assays) |
| AET373 / SAD4064 | *comP*<sup>T129C</sup>, *pilT*<sup>D2K,D53R,K136A</sup> *pilU*<sup>K351D</sup> | Fig 5b | *A. baylyi* natural transformation assays | *pilT*<sup>WA,D2K,D53R</sup>, *pilU*<sup>K351D</sup> (*A. baylyi* NT assays) |
| AET369 / SAD4065 | E7946 Sm<sup>R</sup>, *P*<sub>tac</sub>-*tfoX*, Δ*luxO*, *pilA*<sup>S67C</sup>, Δ*pilT*::Smx<sup>R</sup>, Δ*lacZ*::Spec<sup>R</sup>-*P*<sub>pilT</sub>-*mCherry*-L5-*pilT* | Fig S2 | Fluorescence microscopy foci quantification | *mCherry-pilT*<sup>+</sup>, *pilU*<sup>+</sup> |
| AET367 / SAD4066 | E7946 Sm<sup>R</sup>, *P*<sub>tac</sub>-*tfoX*, Δ*luxO*, *pilA*<sup>S67C</sup>, Δ*pilT*::Smx<sup>R</sup> *mCherry*-L5-*pilU*, Δ*lacZ*::Spec<sup>R</sup>-*P*<sub>pilT</sub>-*pilT* | Fig S2; | Fluorescence microscopy foci quantification | *mCherry-pilU*<sup>+</sup>, *pilT*<sup>+</sup> |
| AET395 / SAD4109 | E7946 Sm<sup>R</sup>, *P*<sub>tac</sub>-*tfoX*, Δ*luxO*, *pilA*<sup>S67C</sup>, Δ*pilT*::Smx<sup>R</sup> *pilU*<sup>K134A</sup>, Δ*lacZ*::Spec<sup>R</sup>-*P*<sub>pilT</sub>-*mCherry*-L5-*pilT*<sup>K136A</sup>, Δ*MSHA*::Carb<sup>R</sup> | Fig S3a | Fluorescence motor and competence pilus colocalization | *mCherry-pilT*<sup>WA</sup>, *pilU*<sup>WA</sup>, Δ*MSHA* |
| AET185 / SAD4110 | E7946 Sm<sup>R</sup>, *P*<sub>tac</sub>-*tfoX*, Δ*luxO*, *pilA*<sup>S67C</sup>, Δ*pilT*::Smx<sup>R</sup> *mCherry*-L5-*pilU*<sup>K134A</sup>, Δ*lacZ*::Spec<sup>R</sup>-*P*<sub>pilT</sub>-*pilT*<sup>K136A</sup>, Δ*MSHA*::Carb<sup>R</sup> | Fig S3a | Fluorescence motor and competence pilus colocalization | *mCherry-pilU*<sup>WA</sup>, *pilT*<sup>WA</sup>, Δ*MSHA* |
| AET484 / SAD4111 | E7946 Sm<sup>R</sup>, *P*<sub>tac</sub>-*tfoX*, Δ*luxO*, *pilA*<sup>S67C</sup>, Δ*pilT*::Smx<sup>R</sup> *pilU*<sup>K134A</sup>, Δ*lacZ*::Spec<sup>R</sup>-*P*<sub>pilT</sub>-*mCherry*-L5-*pilT*<sup>K136A</sup>, Δ*MSHA*::Carb<sup>R</sup>, *msfGFP-pilQ* | Fig S3b-c | Fluorescence motor and PilQ colocalization quantification | *mCherry-pilT*<sup>WA</sup>, *pilU*<sup>WA</sup>, *msfGFP-pilQ*, Δ*MSHA* |
| AET483 / SAD4112 | E7946 Sm<sup>R</sup>, *P*<sub>tac</sub>-*tfoX*, Δ*luxO*, *pilA*<sup>S67C</sup>, Δ*pilT*::Smx<sup>R</sup> *mCherry*-L5-*pilU*<sup>K134A</sup>, Δ*lacZ*::Spec<sup>R</sup>-*P*<sub>pilT</sub>-*pilT*<sup>K136A</sup>, Δ*MSHA*::Carb<sup>R</sup>, *msfGFP-pilQ* | Fig S3b-c | Fluorescence motor and PilQ colocalization quantification | *mCherry-pilU*<sup>WA</sup>, *pilT*<sup>WA</sup>, *msfGFP-pilQ*, Δ*MSHA* |
| AET208 / SAD4067 | E7946 Sm<sup>R</sup>, *P*<sub>tac</sub>-*tfoX*, Δ*luxO*, *pilA*<sup>S67C</sup>, Δ*pilTU*::Tm<sup>R</sup>, Δ*lacZ*::Spec<sup>R</sup>-*P*<sub>pilT</sub>-*pilT*<sup>R35E</sup> | Fig S6a-b | Transformation, pilus labeling, and cellular aggregation assays | *pilT*<sup>R35E</sup> |
| AET209 / SAD4068 | E7946 Sm<sup>R</sup>, *P*<sub>tac</sub>-*tfoX*, Δ*luxO*, *pilA*<sup>S67C</sup>, Δ*pilTU*::Tm<sup>R</sup>, Δ*lacZ*::Spec<sup>R</sup>-*P*<sub>pilT</sub>-*pilT*<sup>K36E</sup> | Fig S6a-b | Transformation, pilus labeling, and cellular aggregation assays | *pilT*<sup>K36E</sup> |
| AET397 /<br>SAD4069 | E7946 Sm<sup>R</sup>, P<sub>tac</sub>-tfoX, ΔluxO,<br>pilA<sup>S67C</sup>, ΔpilTU::Tm<sup>R</sup>,<br>ΔlacZ::Spec<sup>R</sup>-P<sub>pilT</sub>-pilT<sup>D2K,E53R</sup> | Fig S6a-b | Transformation, pilus<br>labeling, and cellular<br>aggregation assays | pilT<sup>D2K,E53R</sup> |
| AET408 /<br>SAD4113 | E7946 Sm<sup>R</sup>, P<sub>tac</sub>-tfoX, ΔluxO,<br>pilA<sup>S67C</sup>, ΔpilT::Smx<sup>R</sup> 3xFLAG-<br>L5-pilU, ΔlacZ::Spec<sup>R</sup>-P<sub>pilT</sub>-<br>pilT<sup>K136A</sup> | Fig S7a | Western blots | pilT<sup>WA</sup> 3xFLAG-<br>pilU<sup>+</sup> |
| AET466 /<br>SAD4114 | E7946 Sm<sup>R</sup>, P<sub>tac</sub>-tfoX, ΔluxO,<br>pilA<sup>S67C</sup>, ΔpilT::Smx<sup>R</sup> 3xFLAG-L5-pilU<sup>K134A</sup>, ΔlacZ::Spec<sup>R</sup>-P<sub>pilT</sub>-<br>pilT<sup>K136A</sup> | Fig S7a | Western blots | pilT<sup>WA</sup> 3xFLAG-pilU<sup>K134A</sup> |
| AET468 /<br>SAD4115 | E7946 Sm<sup>R</sup>, P<sub>tac</sub>-tfoX, ΔluxO,<br>pilA<sup>S67C</sup>, ΔpilT::Smx<sup>R</sup> 3xFLAG-L5-pilU<sup>D366K</sup>, ΔlacZ::Spec<sup>R</sup>-P<sub>pilT</sub>-<br>pilT<sup>K136A</sup> | Fig S7a | Western blots | pilT<sup>WA</sup> 3xFLAG-pilU<sup>D366K</sup> |
| AET477 /<br>SAD4116 | E7946 Sm<sup>R</sup>, P<sub>tac</sub>-tfoX, ΔluxO,<br>pilA<sup>S67C</sup>, ΔpilT::Smx<sup>R</sup> 3xFLAG-L5-pilU<sup>E368K</sup>, ΔlacZ::Spec<sup>R</sup>-P<sub>pilT</sub>-<br>pilT<sup>K136A</sup> | Fig S7a | Western blots | pilT<sup>WA</sup> 3xFLAG-pilU<sup>E368K</sup> |
| AET480 /<br>SAD4117 | E7946 Sm<sup>R</sup>, P<sub>tac</sub>-tfoX, ΔluxO,<br>pilA<sup>S67C</sup>, ΔpilT::Smx<sup>R</sup> 3xFLAG-L5-pilU<sup>K348D</sup>, ΔlacZ::Spec<sup>R</sup>-P<sub>pilT</sub>-<br>pilT<sup>K136A</sup> | Fig S7a | Western blots | pilT<sup>WA</sup> 3xFLAG-pilU<sup>K348D</sup> |
| AET481 /<br>SAD4118 | E7946 Sm<sup>R</sup>, P<sub>tac</sub>-tfoX, ΔluxO,<br>pilA<sup>S67C</sup>, ΔpilT::Smx<sup>R</sup> 3xFLAG-L5-pilU<sup>D366K,E368K</sup>, ΔlacZ::Spec<sup>R</sup>-<br>P<sub>pilT</sub>-pilT<sup>K136A</sup> | Fig S7a | Western blots | pilT<sup>WA</sup> 3xFLAG-pilU<sup>D366K,E368K</sup> |
| AET487 /<br>SAD4119 | comP<sup>T129C</sup>, pilT<sup>K136A</sup> 3xFLAG-<br>pilU | Fig S7b | Western blots | pilT<sup>WA</sup> 3xFLAG-<br>pilU<sup>+</sup> (A. baylyi) |
| AET492 /<br>SAD4120 | comP<sup>T129C</sup>, pilT<sup>K136A</sup> 3xFLAG-pilU<sup>K351D</sup> | Fig S7b | Western blots | pilT<sup>WA</sup> 3xFLAG-pilU<sup>K351D</sup><br>(A. baylyi) |

**Table S2.**
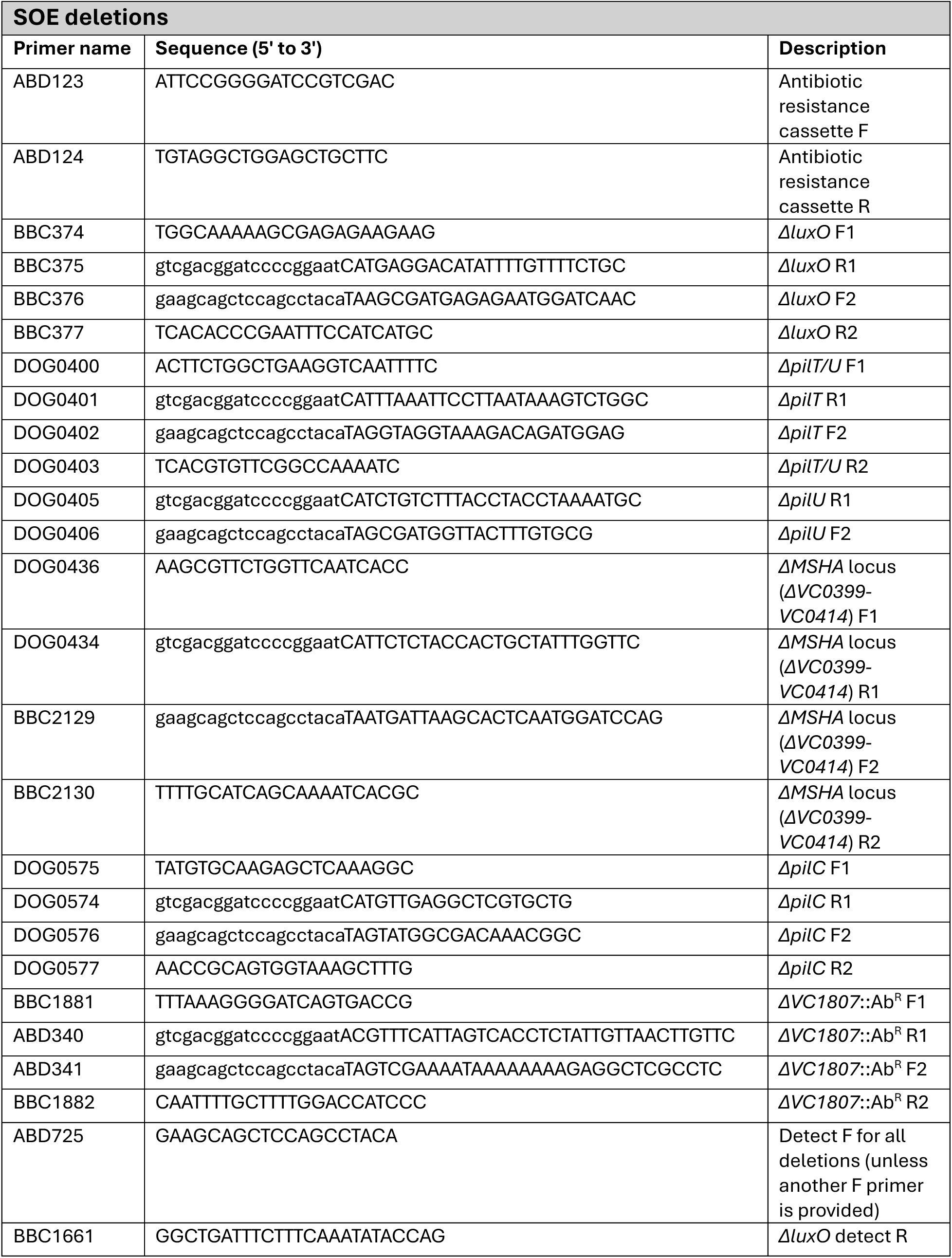
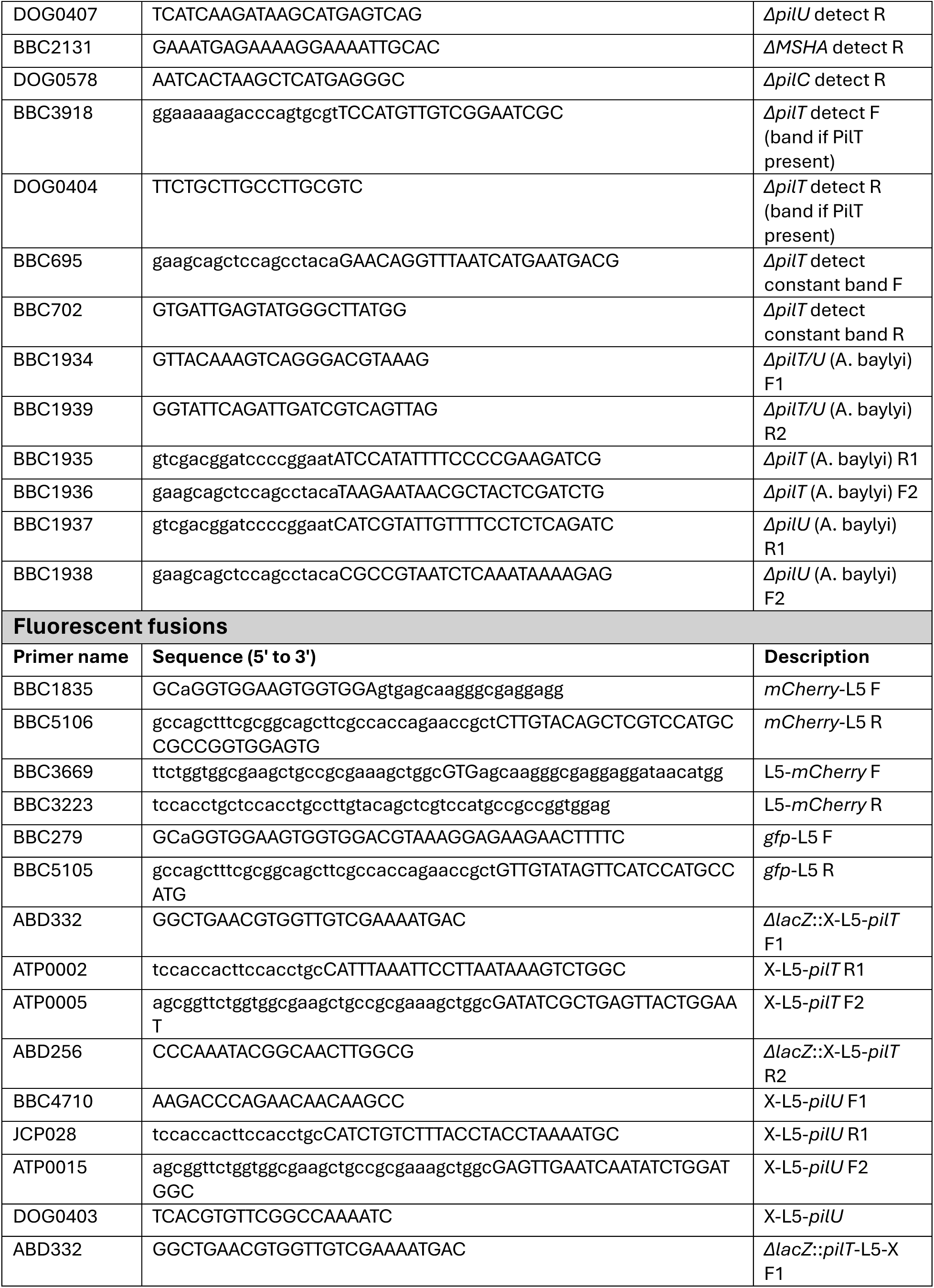
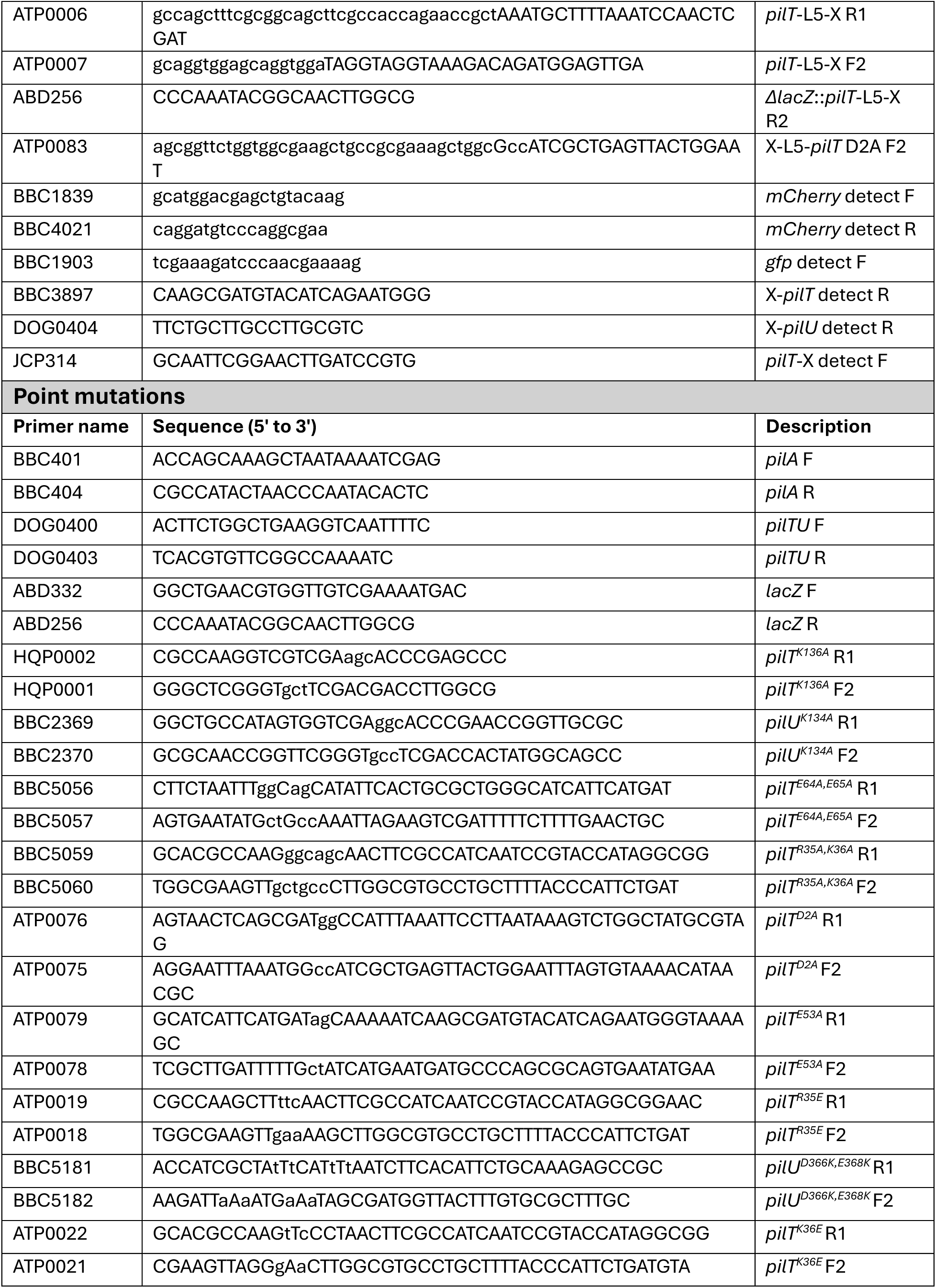
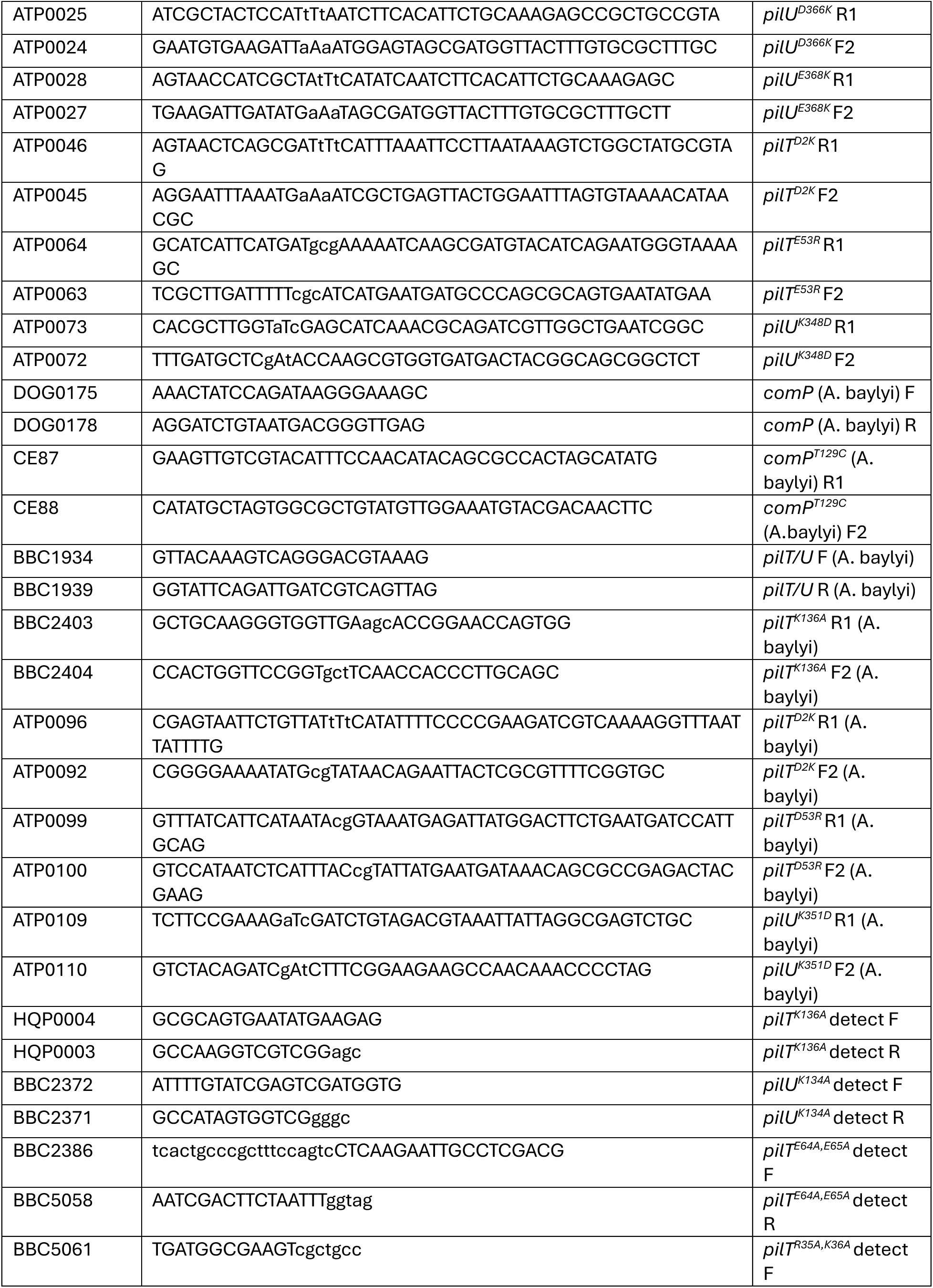
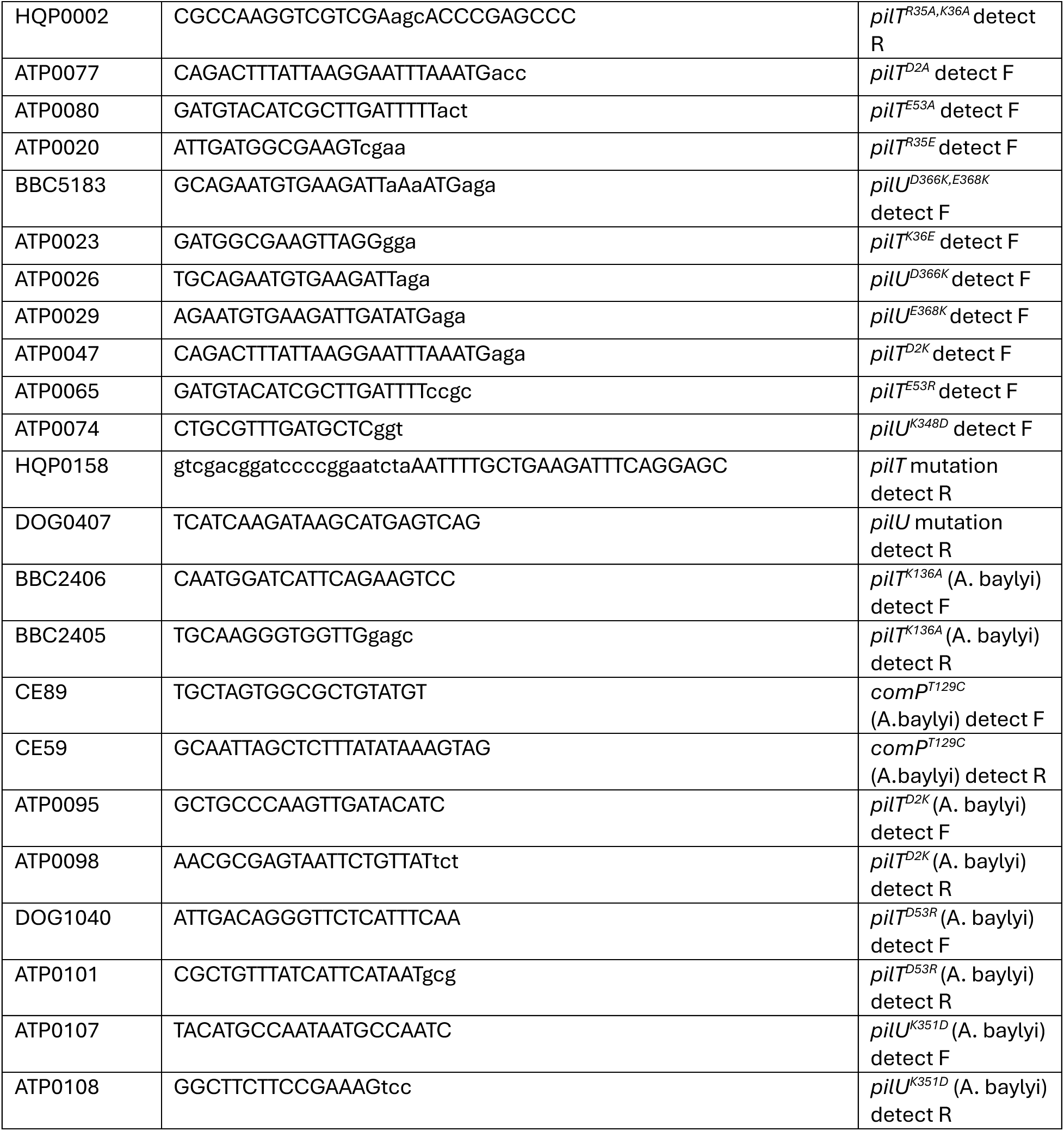
Primers used in this study.

**Movie S1.** Representative movie of the montage shown in Fig S2a. mCherry-PilT^+^ does not form peripheral foci.

**Movie S2.** Representative movie of the montage shown in Fig S2a. mCherry-PilU^+^ does not form peripheral foci.

**Movie S3.** Representative movie of the montage shown in Fig 1b. Motor localization of mCherry-PilT^WA^ shows fluorescent foci at the cell poles.

**Movie S4.** Representative movie of the montage shown in Fig 1b. Motor localization of mCherry-PilU^WA^ foci shows fluorescent foci at the cell poles.

**Movie S5.** Representative movie of the montage from Fig S3a shows mCherry-PilT^WA^ and AF488-mal labeled pilus colocalization.

**Movie S6.** Representative movie of the montage from Fig S3a shows mCherry-PilU^WA^ and AF488-mal labeled pilus colocalization.

**Movie S7.** Representative movie of the montage from Fig S3b shows mCherry-PilT^WA^ and msfGFP-PilQ colocalization.

**Movie S8.** Representative movie of the montage from Fig S3b shows mCherry-PilU^WA^ and msfGFP-PilQ colocalization.

**Movie S9.** Representative movie of the montage from Fig 1e showing GFP-PilU^WA^ and PilT^WA^- mCherry colocalization.

**Movie S10**. Molecular dynamics simulation of the AF3 PilT-PilU model. A 100-ns span of the simulation is shown for PilT^R35^-PilU^D366,E368^.

**Movie S11**. Molecular dynamics simulation of the AF3 PilT-PilU model. A 100-ns span of the simulation is shown for PilT^R35^-PilU^I365,M367^.

**Movie S12**. Molecular dynamics simulation of the AF3 PilT-PilU model. A 100-ns span of the simulation is shown for PilT^K36^-PilU^D366,E368^.

**Movie S13**. Molecular dynamics simulation of the AF3 PilT-PilU model. A 100-ns span of the simulation is shown for PilT^D2,E53^-PilU^K348^.

**Dataset S1**. Summary statistics and a comprehensive list of statistical comparisons for all quantitative data in the manuscript.

